# Environmental stiffness restores mechanical homeostasis in vimentin-depleted cells

**DOI:** 10.1101/2023.05.23.541876

**Authors:** Janine Grolleman, Nicole C.A. van Engeland, Minahil Raza, Sepinoud Azimi Rasthi, Vito Conte, Cecilia M. Sahlgren, Carlijn V.C. Bouten

## Abstract

Recent experimental evidence indicates a role for the intermediate filament vimentin in regulating cellular mechanical homeostasis, but its precise contribution remains to be discovered. Mechanical homeostasis requires a balanced bi-directional interplay between the cell’s microenvironment and the cellular morphological and mechanical state – this balance being regulated via processes of mechanotransduction and mechanoresponse, commonly referred to as mechanoreciprocity. Here, we systematically analyze vimentin-expressing and vimentin-depleted cells in a swatch of *in vitro* cellular microenvironments varying in stiffness and/or ECM density. We find that vimentin-expressing cells maintain mechanical homeostasis by adapting cellular morphology and mechanics to micromechanical changes in the microenvironment. However, vimentin-depleted cells lose this mechanoresponse ability on short timescales, only to reacquire it on longer time scales. Indeed, we find that the morphology and mechanics of vimentin-depleted cell in stiffened microenvironmental conditions can get restored to the homeostatic levels of vimentin-expressing cells. Additionally, we observed vimentin-depleted cells increasing collagen matrix synthesis and its crosslinking, a phenomenon which is known to increase matrix stiffness, and which we now hypothesize to be a cellular compensation mechanism for the loss of vimentin. Taken together, our findings provide further insight in the regulating role of intermediate filament vimentin in mediating mechanoreciprocity and mechanical homeostasis.

## Introduction

Mechanical force controls fundamental processes in health and disease^1^. For a tissue to be functional, cells within the tissue need to establish, maintain, and restore a preferred morphological and mechanical state, a phenomenon referred to as mechanical homeostasis^2^. Loss of mechanical homeostasis is associated with the onset and progression of pathologies, ranging from cardiovascular diseases such as cardiomyopathy^3^ and aneurysms^4^ to cancer^5^. The regulation of mechanical homeostasis requires fine-tuned mechanoreciprocity, which is defined as the dynamic bi-directional mechanical interplay between a cell and its microenvironment rich in extracellular matrix (ECM)^6,7^. Mechanoreciprocity requires mechanical signals from the cellular environment to be sensed by the cells and converted into biomechanical and biochemical signals in the nucleus (mechanotransduction). This process triggers a response from the cell at the mechanical level (mechanoresponse), which in turn leads to adaptation of the cell’s morphological and mechanical (morpho-mechanical) state as well as to the synthesis and remodeling of the ECM^8^.

The complex regulation of the continuous adaptive remodeling process of mechanoreciprocity, which operates over different timescales^6^, is known to involve the cytoskeleton^9,10^. The cytoskeleton is a dynamic network composed of actin, microtubules, and intermediate filaments (IFs), and is important for transmitting mechanical signals from the ECM to the nucleus^11^. The cytoskeleton also governs the morpho-mechanical state of a cell, and extensive experimental evidence has elucidated the roles of stretch-resistant actin and compression-resistant microtubules in maintaining cellular mechanical homeostasis^12^. Yet, little is still known about the contribution of IFs to the mechanical homeostatic balance.

The IF protein vimentin is involved in many processes driven by cell mechanics such as wound healing, angiogenesis, flow-induced arterial remodeling, and closure of the ductus arteriosus (reviewed in ref 13). Vimentin is highly expressed in mesenchymal cells, including fibroblasts, endothelial cells (ECs), and vascular smooth muscle cells (VSMCs)^14^, where it appears as a cage-like network surrounding the nucleus and is crucial in regulating nuclear morphology^15^. The vimentin network extends to the cell cortex, including protrusions, where it forms an interpenetrating network with the actin cytoskeleton^16^. Both vimentin and actin associate with focal adhesions (FAs), the protein complexes that mechanically couple the cytoskeleton to the ECM at the cell membrane^17,18^. FAs dynamics is known to be controlled by vimentin^19–21^ and is highly dependent on the mechanical characteristics of the ECM, including ECM stiffness and density^22^. Recent studies have been highlighting a regulatory role for vimentin in the synthesis and remodeling of the ECM *in vivo*, as increased ECM production and tissue stiffness is observed in vimentin-depleted mice^23,24^. Thus, we hypothesize that vimentin regulates mechanoreciprocity, but *how* vimentin contributes to mechanical homeostasis is not known (Supplementary Fig. S1).

In this study, we systematically investigate the role of vimentin in mechanical homeostasis by quantifying how vimentin depletion affects ECM expression and the morpho-mechanical state of cells from arterial tissues cultured in a swatch of mechanical microenvironmental conditions varying in stiffness and ECM density. We find that vimentin is essential for cells to restore mechanical homeostasis on shorter timescales by initiating an adaptive mechanical response to changes in stiffness and ECM density of the cell’s microenvironment. Our data shows that vimentin depletion in cells initially disrupts their mechanoresponsive ability and then triggers an increase in their expression of ECM components and collagen crosslinkers on a longer timescale. Surprisingly, vimentin-depleted cells are able to restore their morphology and mechanics to the homeostatic levels of vimentin-expressing cells, provided that the stiffness of these cells’ microenvironment is sufficiently increased independently of ECM density.

## Results

### Vimentin regulates ECM production and remodeling

Previous research has shown that loss of vimentin results in increased ECM production and remodeling *in vivo*^23,24^. To better understand the changes in ECM turnover due to loss of vimentin we cultured vimentin-wildtype (VimWT) and vimentin-knockout (VimKO) mouse embryonic fibroblasts (MEFs) on collagen type I coated polyacrylamide (PAA) hydrogel substrates having cardiovascular physiological stiffness of 12 kPa^25,26^. Vimentin knockout was verified on both protein (Fig. 1A,B) and gene level (Fig. 1C). Next, we stained VimWT and VimKO cells with an antibody against collagen type III to discriminate from the collagen type I coating. After 72 hours of cell culture on the hydrogel, VimKO cells expressed a net increase in collagen type III as compared to VimWT cells (Fig. 1D), which is in line with the *in vivo* observations^23,24^. To refine the temporal effect and the balance in synthesis and remodeling of this finding, we analyzed the relative mRNA expression of ECM synthesis and remodeling genes in VimKO cells and VimWT cells after 24 and 72 hours. The loss of vimentin significantly increased mRNA levels of *Col3a1* and *Col1a1* after 24 hours of cell culture (Fig. 1E). A similar vimentin depletion related increase in ECM synthesis was observed in other vascular cell types, endothelial (Supplementary Fig. S2) and vascular smooth muscle cells (Supplementary Fig. S3), which highlight that this phenomenon is a cell generic response. In addition, we observed an increase in the expression of the collagen crosslinking gene *Lox* (Fig. 1F) and the matrix remodeling genes *Mmp2* and *Timp2* (Supplementary Fig. S4A). After 72 hours of culture on the 12 kPa substrate, the increase in *Col3a1* and *Lox* normalized to the expression levels of VimWT cells, while the expression of *Col1a1, Mmp2* and *Timp2* remained increased. This suggests that there is an initial boost of collagen type III synthesis and collagen crosslinking, and prolonged synthesis of collagen type I and matrix remodeling.

**Figure 1.**
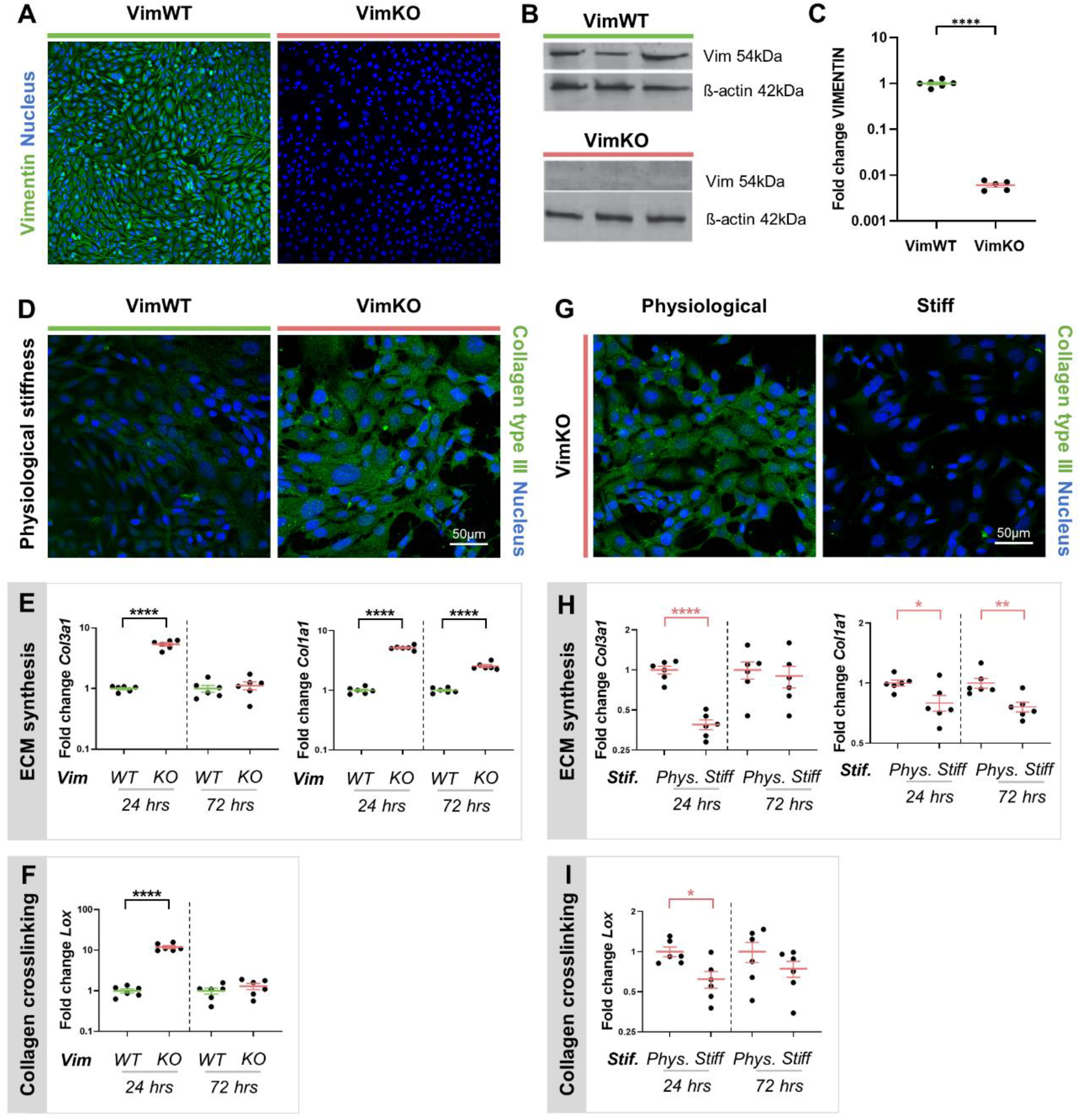
Increased substrate stiffness compensates for vimentin depletion altered ECM synthesis and crosslinking. Verification of vimentin knockout on protein level using immunofluorescence (IF) staining (A) and Western Blotting (B), and on gene level using q-PCR with respect to Gapdh expression and normalized to VimWT expression levels (C). (D-F) VimWT and VimKO cells cultured on PAA gels with physiological substrate stiffness (12 kPa). Collagen type III protein expression after 72 hours by IF staining (D) and gene expression after 24 and 72 hours by q-PCR of ECM synthesis gene Col3a1 and Col1a1 (E), and collagen crosslinking gene Lox (F) with respect to Gapdh expression and normalized to VimWT expression levels. (G-I) VimKO cells cultured on PAA gels with physiological substrate stiffness (Phys.) and on a stiff substrate (glass). Collagen type III protein expression after 72 hours by IF staining (G) and gene expression after 24 and 72 hours of ECM synthesis genes Col3a1 and Col1a1 (H), and collagen crosslinking gene Lox (I). N=3 for WB and N=6 for IF staining and q-PCR. Data is represented as mean±SEM. An unpaired t-test was used for statistical analysis. Denotations: **** p<0.0001; *** p<0.001; ** p<0.01; * p<0.05; · p<0.1.

Apart from increased ECM production, *in vivo* studies showed arterial stiffening as a result of vimentin depletion. To investigate the effect of changes in stiffness of the microenvironment, we cultured VimKO cells on physiological substrate stiffness and high substrate stiffness (glass) to mimic arterial stiffening and compared the production of ECM components and regulators. Contrary to VimKO cells on physiological stiffness, VimKO cells cultured on stiff substrates showed reduced collagen type III protein levels (Fig. 1G). Further analysis demonstrated a decrease in the expression of matrix genes *Col3a1, Col1a1* (Fig. 1H), *Lox* (Fig. 1I), *Mmp2*, and *Timp2* (Supplementary Fig. S4B) after 24 hours of VimKO cell culture on stiff substrates compared to physiological stiffness. After 72 hours of cell culture, VimKO expression levels on stiff substrates normalized to the expression levels of VimKO cells on physiological stiffness except for ECM synthesis gene *Col1a1 and* remodeling gene *Timp2*. Taken together, these data show that vimentin mediates ECM synthesis, crosslinking and remodeling depending on environmental stiffness and suggest that vimentin-depleted cells increase their extracellular matrix production unless they are cultured on substrates with increased stiffness.

### Vimentin-depleted cells adopt a differential morpho-mechanical phenotype

The observed changes in the ECM composition (stiffness and ECM density) prompted us to study how these mechanical alterations within the microenvironment of the cell affect the morpho-mechanical state of the cell. To this end, we cultured VimWT and VimKO cells on substrates with physiological (E_P_) or high (E_H_) substrate stiffness and with physiological (C_P_) or high (C_H_) substrate (collagen type I) ECM density (Fig. 2A) and studied the morpho-mechanical adaptation response after 24 hours. We stained for cytoskeletal protein F-actin and acquired top- and side view images (Fig. 2B).

**Figure 2.**
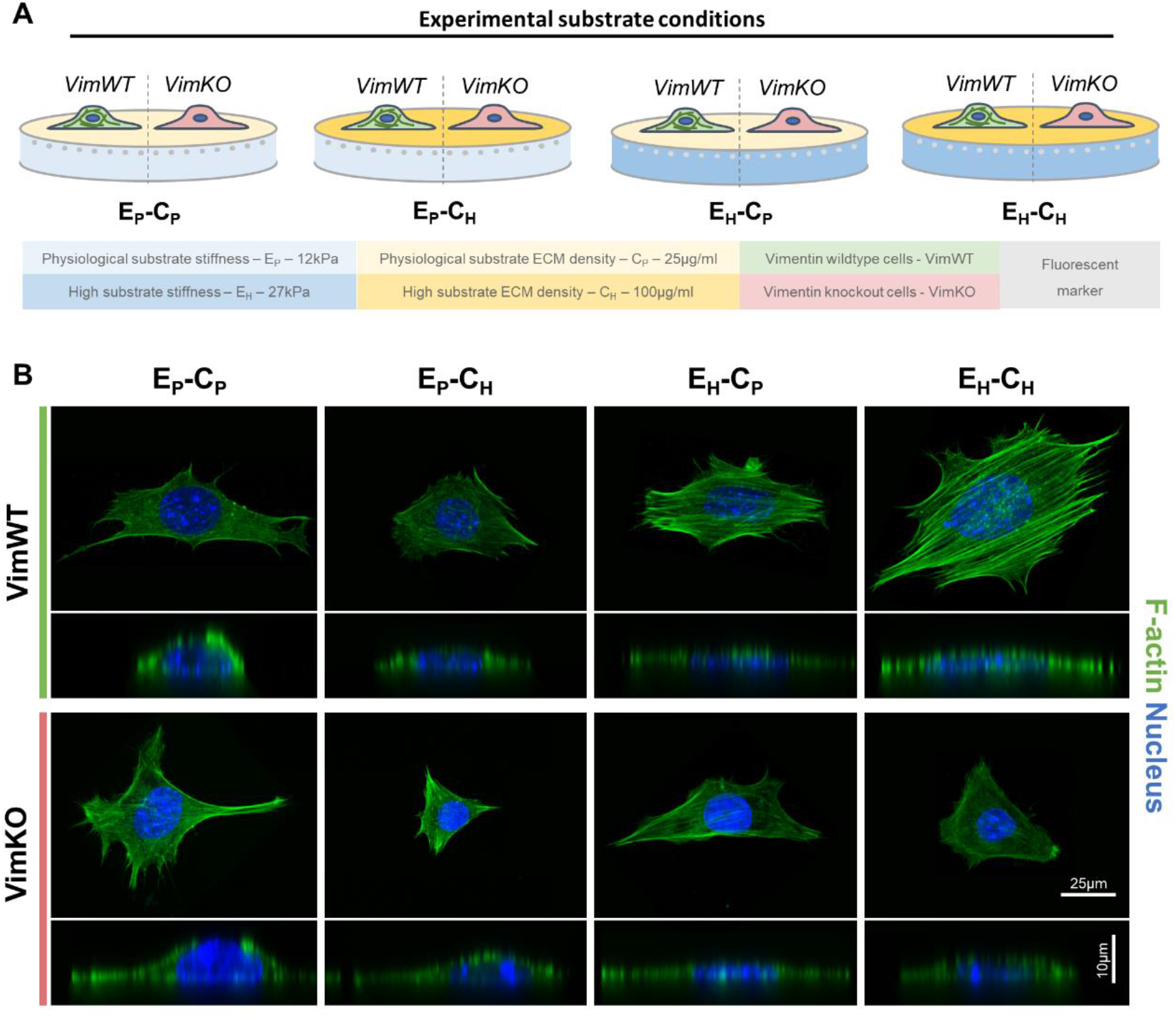
Substrate conditions alter morpho-mechanical state of the cell. (A) Schematic representation of VimWT and. VimKO cells cultured on different substrate conditions: physiological (E_P_) vs stiff (E_H_) substrate stiffness and physiological (C_P_) vs high (C_H_) substrate ECM density. VimWT and VimKO cells were cultured on the different substrate conditions for 24 hours. (B) Top and side confocal images of VimWT and VimKO cells cultured on different substrate conditions for 24 hours stained for F-actin and DAPI (N=6).

To observe differences in the morpho-mechanical state of the cell, we provide a quantitative measure of the morpho-mechanical state of the cell. Briefly, the morphological state was quantified by nuclear and cellular morphology, whereas the mechanical state was quantified by FA formation and cellular traction forces (Fig. 3A). To characterize cellular morphology, we quantified cell area and height. Both VimWT and VimKO cells showed an increasing trend in cell area to increased substrate stiffness (Fig. 3B) which is in agreement with the paradigm of increased cell spreading on stiffer substrates^27–30^. In line with this, both cell types decreased in height when cultured on hydrogels with either high substrate ECM density on physiological stiffness or high substrate stiffness with physiological ECM density, whereas combined high substrate stiffness and high substrate ECM density did not significantly affect cell height (Fig. 3C). Next, we examined if vimentin affects nuclear morphology by acquiring top- and side view images of the nucleus (Supplementary Fig. S5A) and quantified nucleus area and height. Quantification of the area of the nucleus showed a reduction in the area in VimKO cells compared to VimWT cells (Supplementary Fig. S5B). VimWT cells did not adapt the height of the nucleus to changes in the microenvironment whereas VimKO cells significantly increased nuclear height on hydrogels with increased ECM density on physiological stiffness or increased substrate stiffness with physiological ECM density as compared to VimWT cells (Fig. 3D). While our data on nuclear morphology is in line with previous research showing that vimentin depletion results in rounder nuclei^15,31^, we found that cellular morphology is not affected by the presence of vimentin, most likely since vimentin is highly localized in the core of the cytoplasm and to a lesser degree at the cell periphery^32^.

**Figure 3.**
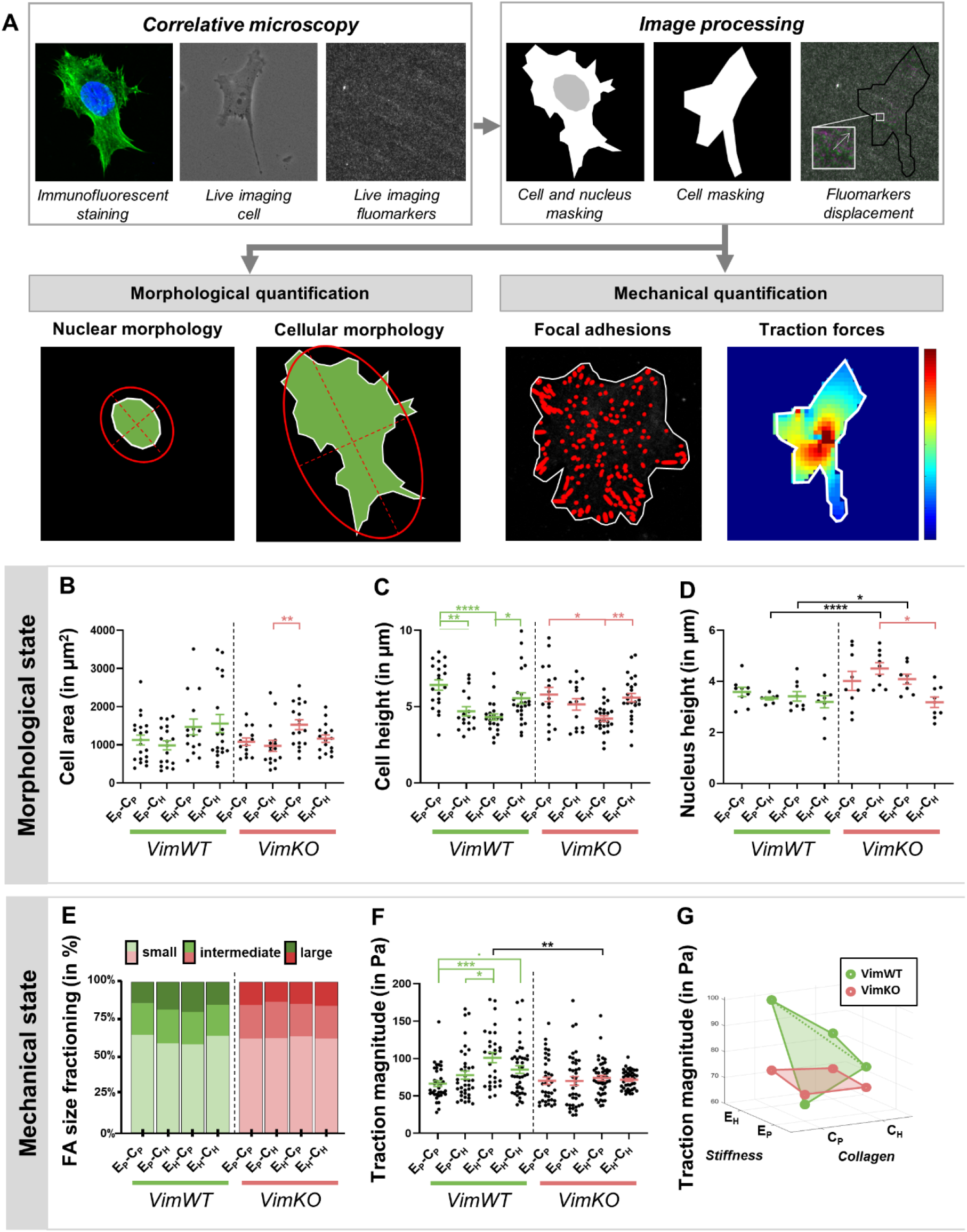
Vimentin depleted cells lose the ability to mechanically respond to alterations in the microenvironment after 24 hours. (A) Schematic representation of image quantification workflow (see Materials and Methods section for detailed explanation). (Correlative microscopy) Phase contrast and fluorescence imaging was used to visualize the cell and/or the fluorescent markers (fluomarkers) within the gel. (Image processing) Cell and nucleus were manually masked and fluomarkers displacement with respect to the reference image was processed by PIV (right panel; reference in green and cell contraction in magenta) and converted into a displacement vector (white arrow). (Morphological quantification) Cellular and nuclear morphology are based on the masks and quantification includes mask size (green), mask perimeter (white), ellipse ratio defining shape (red). Height was determined directly from the IF staining images. (Mechanical characterization) The mask is used to only extract data within the cell. For FA quantification, a binary image of detected FAs was produced, ellipses were fitted on the FAs and FA number, size (ellipse ratio) and shape were extracted from this analysis. FA fractioning was based on combined FA number and size. For traction force analysis, displacement vectors were converted into traction vector using the mechanical properties of the PAA gel. Traction magnitude was computed by taking the median magnitude within the mask and mean over the timelapse. (B-D) Morphological state quantification of cell area (B; N=15+), cell height (C; N=15+), and nucleus height (D; N=8+). Each dot represents one cell/nucleus. (E-G) Mechanical state quantification of FAs (E; N=8+) and traction magnitude (F-G; N=31+). (E) Each dot represents on cell averaged over time. (F) Dots represent group mean and are connected in a bending plane for VimWT cell (in green) and a flat plane for VimKO cells (in red). Data is represented as mean±SEM. Kruskall-Wallis followed by Dunn’s multiple comparison was used for statistical analysis between substrate conditions within VimWT cells (displayed in green) and VimKO cells (displayed in red). A Mann-Whitney test was used for statistical analysis between VimWT and VimKO cells on different substrate conditions (displayed in black). Denotations: **** p<0.0001; *** p<0.001; ** p<0.01; * p<0.05; · p<0.1.

As for the mechanical state of the cell we studied the formation of actin stress fibers, myosin II activity, FA formation and tractions forces^33^, which are known contributors to cellular functions including cell morphology, cell migration and ECM organization^34^. Both VimWT and VimKO cells display comparable F-actin organization when cultured on physiological stiffness independently of ECM density (Fig. 2B). However, VimWT cells formed abundant actin stress fibers when cultured on high substrate stiffness independently of ECM density, whereas VimKO cells did not exhibit comparable actin stress fibers to VimWT cells on high substrate stiffness (Fig. 2B). Besides actin organization, we assessed the effect of vimentin depletion on myosin II activity by detection of phosphorylated myosin light chain (pMLC). Independent of the substrate, VimKO cells showed an increased level of pMLC compared to VimWT cells (Supplementary Fig. S5C), which is in agreement with previous findings by Jiu and colleagues^35^. Vimentin depletion also affected FA formation on the different substrates as demonstrated by quantification of the FA size based on a staining for paxillin (Supplementary Fig. S5D). Upon fractioning the FAs into small, intermediate, and large FAs within our analysis, we observed that increased substrate ECM density on physiological stiffness and increased substrate stiffness with physiological ECM density resulted in more large FAs in VimWT cells, but no change was observed in VimKO cells (Fig. 3E). Next, we examined how these changes in the mechanical components of the cell affected cell traction forces using traction force microscopy. On physiological substrate conditions, VimWT and VimKO cells show a similar contractile behavior (Fig. 3F). While VimWT cells increased their traction magnitude to increasing substrate stiffness with physiological ECM density, VimKO cells did not adapt traction magnitude upon changes in the microenvironment (Fig. 3F). A significant decrease in traction magnitude was observed in VimKO cells cultured on high substrate stiffness with physiological ECM density as compared to VimWT cells (Fig. 3F). Interestingly, VimWT cells increased traction magnitude upon increased substrate ECM density on physiological substrate stiffness, whereas they decreased traction magnitude upon increased substrate ECM density on high substrate stiffness (Fig. 3G) indicating that there is an optimal substrate condition at which VimWT cells are able to exert the highest cell tractions on the substrate. A similar trend was observed when traction magnitude was normalized for cell size (Supplementary Fig. S5E) suggesting that increased traction magnitude on high substrate stiffness is not related to an increase in cell area. This data shows that vimentin is essential for stress fiber formation and the formation of larger focal adhesions and consequently increased traction force as a mechanoresponse to changes in the mechanical properties of the microenvironment.

The diversity of the morphological and mechanical response presented by cell with and without vimentin prompted us to systematically study the morpho-mechanical state explored by cells as a response to changes in the microenvironment. To do so, we started by examining whether the mechanical and morphological properties defining the cell’s response to microenvironmental changes were linked through universal relationships or whether they were generally uncorrelated^36^. To that end, we first identified a number of properties that captured the variety of morphological and mechanical responses of the cell to changes in stiffness and ECM density of the microenvironment. Next, we averaged these properties over experimental repeats and grouped them as follows: 1) cellular morphology (i.e. cell area, perimeter, height and shape); 2) nuclear morphology (i.e. nucleus area, perimeter, height and shape); 3) cell mechanics (i.e. traction magnitude, average normal stress and maximum shear stress); 4) cell migration (i.e. migration pace and persistence); and, 5) cell-matrix interactions (i.e. FA number, size and shape). This way we could summarize our data into a *p*x*q* matrix *D* (data not shown), whose *p*-rows represent the cellular microenvironment conditions with and without vimentin (*p*=8) and *q-*columns represent the average morphological and mechanical property of the cells in those environmental conditions (*q*=16). From this matrix *D*, we computed a new *m*x*n* matrix *Z* (*m*=*p*-1=7 and *n*=*q*=16, Fig. 4A). Each element of matrix *Z* represents the Z-score of a specific morpho-mechanical property of the cell as a response to a specific microenvironmental condition with respect to the control condition E_P_-C_P_-VimWT (i.e. vimentin-expressing cells on a substrate having physiological stiffness and physiological ECM density). Thus, a positive Z-score signifies that the mean of a specific physical property of a cell in a specific environmental condition is higher compared to the control situation, while a negative Z-score signifies that it is lower than the control situation. Based on this *Z* matrix, we computed the cross-correlation matrices *C* (Fig. 4B) and *P* (Fig. 4C) which explored similarities between environmental conditions and morpho-mechanical properties respectively. A positive similarity between two environmental conditions signals that cells respond to these environmental conditions with similar morpho-mechanical properties, whereas a positive similarity between two morpho-mechanical properties signals that cells respond highly similar in terms of these properties in all environmental conditions. A negative similarity signifies that those environmental conditions/morpho-mechanical properties highly differ from each other. A similarity close to zero signifies that some environmental conditions/morpho-mechanical properties are similar while some other environmental conditions/morpho-mechanical properties are different. Consequently, we used an unsupervised cluster algorithm^36,37^ to identify clusters with highly correlated environmental conditions (Fig. 4B) and morpho-mechanical properties (Fig. 4C). Using the unsupervised cluster algorithm, we identified two clusters of highly correlated environmental conditions contoured with black lines (Fig. 4B). The first cluster consisted of all VimWT conditions, while the second cluster constituted of all VimKO conditions. The unsupervised clustering algorithm detected two correlated clusters of physical properties (Fig. 4C). The first cluster demonstrated a high correlation between nuclear morphology and cell migration, which is in line with previous research^38^. Within the second cluster, we observed a high correlation between cellular morphology and cell mechanics. We further utilized the unsupervised clustering analysis of environmental conditions and morpho-mechanical properties to reorganize the initial matrix *Z* (Fig. 4D). Nuclear morphology and cell migration were negatively affected in VimKO cells while no strong effect was observed in VimWT cells. For cellular morphology and cell mechanics, VimWT displayed a strong increase in Z-score compared to VimKO cells independently of substrate. This highlights the differential morpho-mechanical state of vimentin-depleted cells in comparison to vimentin-expressing cells. Taken together, this analysis suggests that vimentin is required for cells to adapt to changes in the micromechanical environment within 24 hours. However, mechanoreciprocity occurs at different timescales^6^ which may result in cells being able to restore mechanical homeostasis on a variety of timescales. Thus, we wondered whether vimentin-expressing and vimentin-depleted cells had reached mechanical homeostasis (and thus if differences in their responses were permanent or only transitory).

**Figure 4.**
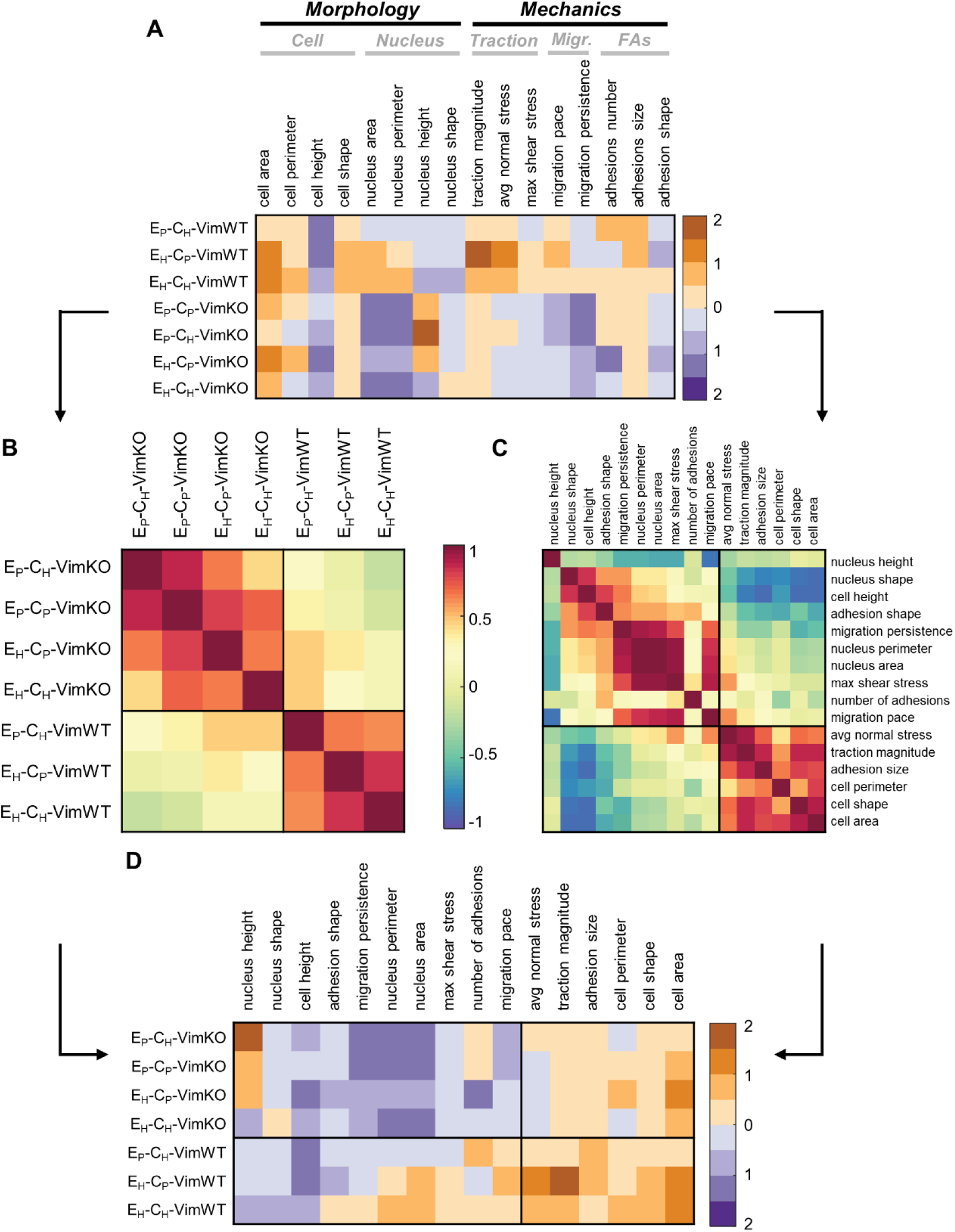
Vimentin depleted cells adopt differential morpho-mechanical phenotype after 24 hours. (A) Matrix *Z* containing Z-Score of morpho-mechanical properties of different microenvironmental conditions with respect to control culture condition E_P_-C_P_-VimWT. (B-C) Ordered and clustered correlation matrices *C* and *P* between environmental conditions (B) and morpho-mechanical properties (C) respectively. (D) Ordered matrix *Z* based on clusters of conditions and properties correlation matrices.

### Vimentin is not necessary for establishing mechanical homeostasis

To test whether cells had reached a homeostatic morpho-mechanical state after 24 hours, we cultured VimWT and VimKO cells on the different substrates (Fig. 2A) for 48 hours and investigated the morphological and mechanical state of the cell. Specifically, we restricted our monitoring of the morpho-mechanical state of the cell at 48 hours to cellular morphology and cell mechanics since these morpho-mechanical properties highly correlated in our similarity analysis at 24 hours (Fig. 4C). For VimWT cells, cell area and cell traction forces at 24 hours (Supplementary Fig. S6A,B) and 48 hours (Supplementary Fig. S6C,D) varied across environmental conditions in a similar fashion indicating that VimWT cells have reached a mechanical homeostatic state already at this timepoint. Surprisingly, VimKO cells showed a similar mechanoresponse as VimWT cells at 48 hours across microenvironmental conditions (Supplementary Fig. S6D) while VimKO cells did not adapt its traction forces to its microenvironment after 24 hours (Supplementary Fig. S6B). This suggests that vimentin-depleted cells need time to respond and restore a mechanical homeostatic state. Nevertheless, VimKO cells cultured on physiological substrate stiffness showed decreased traction magnitude compared to VimWT cells while traction magnitude of VimKO cells on high substrate stiffness with physiological ECM density was comparable to that of VimWT cells (Supplementary Fig. S6D).

To corroborate the different response in time of VimKO cells, we repeated the unsupervised clustering analysis on experimental data sets collected at 24 and 48 hours of cell culture. Cross-correlations of microenvironmental conditions and Z-score reorganization at 24 hours shows that the presence of vimentin determines the morpho-mechanical phenotype of the cell (Fig. 5A,B). When culturing cells for 48 hours on the substrates, the unsupervised cluster analysis again identified two clusters (Fig. 5C,D), these clusters presenting remarkable differences compared to two identified clusters at 24 hours. The data at 48 hours showed that VimKO cells can respond similarly to VimWT cells provided that the stiffness of the microenvironment is high enough, a feature not shown at 24 hours of cell culture. These data highlight that vimentin is not required to establish mechanical homeostasis and indicate that there is a compensation mechanism of vimentin-depleted cells through modulation of matrix stiffness and time to establish a mechanical homeostatic state.

**Figure 5.**
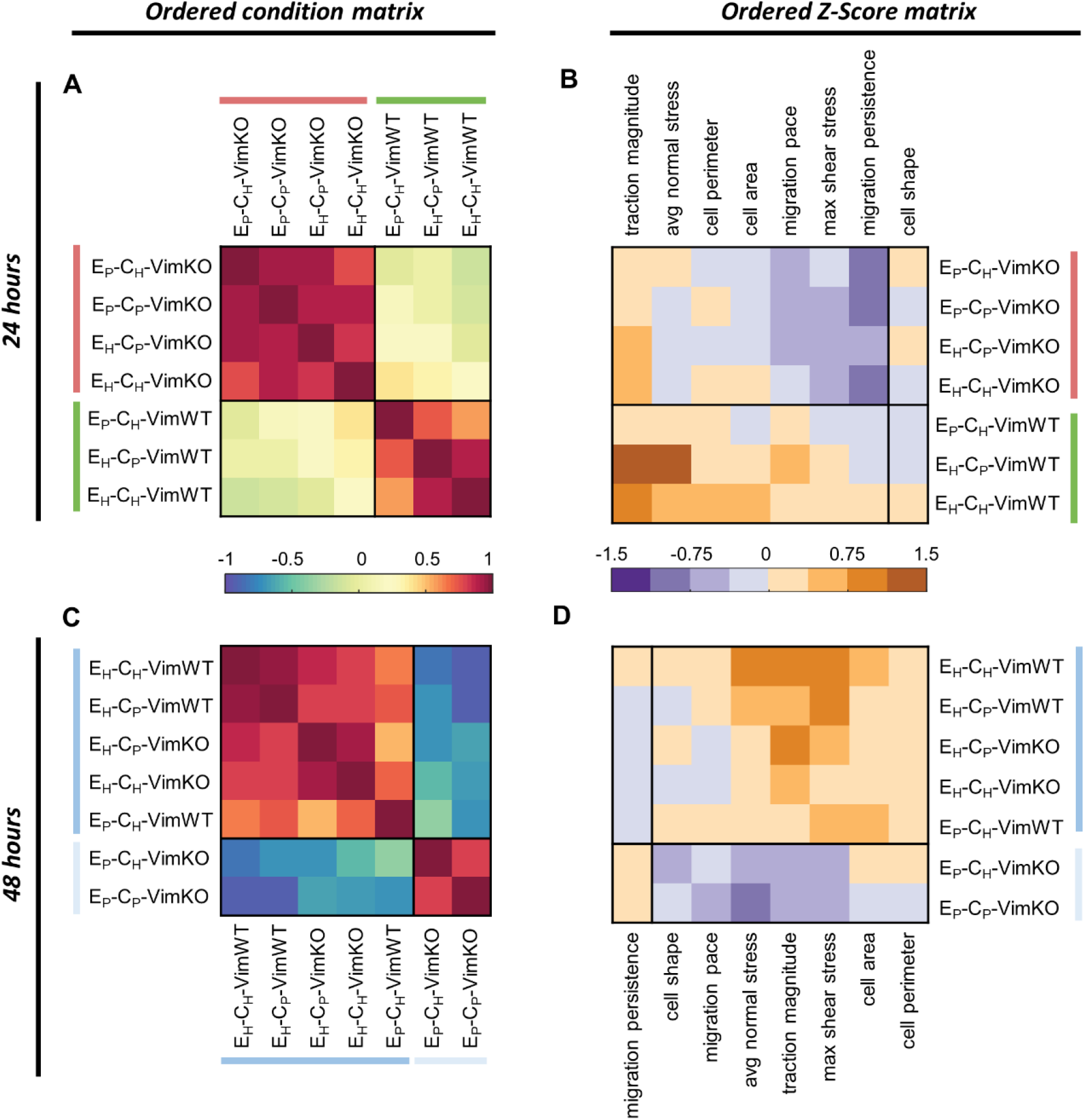
Increased substrate stiffness rescues loss of morpho-mechanical phenotype in vimentin depleted cells after 48 hours. VimWT and VimKO cells were cultured for 24 hours (A-B) or 48 hours (C-D) on the different microenvironmental conditions. Ordered and clustered correlation matrices *C* between environmental conditions (A&C) and *P* between morpho-mechanical properties (data not shown) were produced and used to order matrix *Z* (B&D).

## Discussion

Cellular mechanical homeostasis is established and maintained through the bi-directional mechanical interplay between the cell and the ECM, a dynamic process referred to as mechanoreciprocity. In this process, cells sense and transduce the mechanical stimuli from the ECM (mechanotransduction) and respond mechanically to them by adapting the morpho-mechanical state of the cell as well as the production and remodeling of ECM by the cell (mechanoresponse). Increasing evidence suggests that the IF vimentin contributes to the process of mechanoreciprocity. Hence, alterations in a cell’s IFs may ultimately lead to the disruption of the mechanical homeostatic state of that cell. Here we showed that culturing vimentin-depleted cells on substrates with increased stiffness compensates for the increased expression of ECM and collagen crosslinkers due to loss of vimentin (Fig. 1). Concurrently, increased matrix stiffness – independent of ECM density – also compensates for vimentin depletion in the restoration of the morpho-mechanical phenotype of the cell though on a longer time scale (Fig. 5).

We systematically quantified the morpho-mechanical state of vimentin-expressing and vimentin-depleted MEFs by quantifying cellular and nuclear morphology, cell traction forces, cell migration and FA formation on collagen type I coated PAA hydrogel (substrate matrix) having different stiffness and ECM collagen densities (microenvironment). On substrates having physiological stiffness and ECM density, vimentin-expressing and vimentin-depleted cells behave similarly in terms of cellular and nuclear morphology, cell traction magnitude and FA size (Fig. 3). However, changes in the mechanical properties of the microenvironment trigger different mechanical adaptive responses in vimentin-expressing and vimentin-depleted cells. Differently from vimentin-depleted cells, vimentin-expressing cells were able to adapt their morpho-mechanical state to changes in mechanical properties of the microenvironment (stiffness and/or ECM density). Additionally, vimentin-expressing cells presented an optimal response to substrate stiffness and ECM density in terms of larger FAs, higher tractions and increased cell spreading – both features are lost in vimentin-depleted cells (Fig. 3). A cellular optimal response to changes in the microenvironment has been previously observed in several mechanobiological processes including cell spreading^28,29,39,40^, cell traction^40,41^, cell migration^39,40,42–44^, and YAP localization^45^. This optimal response may rely on the cell’s ability to form FAs, a dynamic process that is highly dependent on both substrate stiffness and ECM density and affects the mechanosensing capabilities of a cell. ECM density determines the number of FAs per cell^45^, while substrate stiffness in combination with the number of FAs determine the actomyosin contractility transmitted to the FAs, which in turn stimulates both growth and stabilization of these FAs^46^. FAs have been shown to stabilize faster on stiffer substrates due to a lower disassembly rate^47,48^. Upon increasing stress on FAs, cells form actin stress fibers that results in increased force loading of FAs – this process acts as a feedback loop in the growth and stabilization of FAs^48^. Through this loop, vimentin-expressing cells can sense and respond to changes in the mechanical properties of their microenvironment by adapting their morpho-mechanical state to restore mechanical homeostasis. Vimentin-depleted cells, instead, lose this mechanical property along with the IFs (Fig. 3).

Previous research investigated the effect of vimentin expression on FAs in terms of their expression levels^23,49,50^, numbers^21,51,52^, sizes^20,21,49,52–54^, and dynamics^19–21,52–54^ while hinting at a regulatory role for vimentin in the formation of FAs. Additionally, Ostrowska-Podhorodecka and colleagues have proposed that vimentin acts as an adaptor protein for FA proteins^49^. By suggesting that vimentin may likely regulate both growth and stabilization of FAs, these studies also implied that loss of vimentin may affect the mechanosensing capabilities of a cell. In this study, we observed no changes in FA size, impaired stress fiber formation, and no response in cell traction force in vimentin-depleted cells upon mechanical changes in the microenvironment (Fig. 2 and 3). This result highlights that the early morpho-mechanical phenotype of MEF cells is determined by vimentin expression (Fig. 4). Vimentin-expressing cells reach a mechanical homeostatic state earlier (24 hours) than vimentin-depleted cells and show no further changes in their morpho-mechanical features with time (Fig. 5). Instead, vimentin-depleted cells reach this state at a longer timescale (48 hours). Vimentin-depleted cells cultured on a stiff substrate independently of ECM density for 48 hours adopt a phenotype comparable to that of vimentin-expressing cells independently of substrate condition (Fig. 5). Thus, our data demonstrates that vimentin-expressing and vimentin-depleted cells can restore mechanical homeostasis at different timescales. In that respect, it is worth noticing that production and remodeling of ECM is a process occurring at different timescales compared to the adaptation of the morpho-mechanical state of a cell. Indeed, previous research has provided evidence for a regulating role of vimentin in stabilization of collagen mRNAs^55,56^, collagen fiber alignment^57^ and ECM production and remodelling^23,24^. Our data on physiological substrate stiffness show that vimentin depletion results in increased ECM synthesis and collagen crosslinking (Fig. 1), processes that have been correlated with a local increase in environmental stiffness^58,59^. Also, culturing vimentin-depleted cells on stiffer substrates decreases ECM production and collagen crosslinking in comparison to physiological substrate stiffness (Fig. 1).

Concluding, we have further elucidated the role that vimentin plays in mediating processes of cellular mechanoreciprocity and, consequently, mechanical homeostasis. Indeed, while the restoration of mechanical homeostasis for cells is not hindered on longer timescales, IF vimentin provides cells with a fast mechanical adaptive response to changes in the mechanical properties of the microenvironment. In that respect, our results point towards the existence of a cellular mechanism that enables vimentin-depleted cells to compensate their loss of vimentin in restoring mechanical homeostasis, even if only on a slower timescale. We speculate that this compensatory mechanism consists in vimentin-depleted cells increasing their matrix production and collagen crosslinking capacity (as observed in Fig. 1), which will ultimately result in the stiffening of the matrix surrounding the cells. With time, this compensatory matrix stiffening may allow vimentin-depleted cells to restore the homeostatic morpho-mechanical phenotype characteristic of vimentin-expressing cells (as observed Fig. 5).

## Materials and Methods

### Cell culture

Vimentin wildtype (VimWT) and vimentin knockout (VimKO) mouse embryonic fibroblasts (MEFs) were kindly provided by John Eriksson (Åbo Akademi, Finland).

### Polyacrylamide (PAA) substrate preparation

PAA gels with a Young’s modulus of 12 kPa (E_P_) and 27 kPa (E_H_) were prepared on glass bottom well plates (MatTek/CellVis) of microscope glass slides. PAA gels were functionalized using 1.0 mg/ml Sulfo-SANPAH (Pierce) and coated with 25 μg/ml (C_P_) or 100 μg/ml (C_H_) rat tail collagen type I (Corning).

### Immunofluorescence staining

MEFs were fixed in paraformaldehyde (ThermoFisher), permeabilized using Triton-X-100 (Sigma) and blocked in bovine serum albumin (Roche). Primary antibodies used in this study: Collagen type III (ab7778, abcam), Paxillin (ab32084, abcam), phosphorylated myosin light chain (3675S, Cell Signaling) and Vimentin (ab20346, abcam). F-actin, collagen and the nucleus were stained using Phalloidin (Sigma), CNA probe (CNA35-OG488) and DAPI (Merck) respectively. Images were acquired using an epifluorescence (Leice DMi8) or confocal microscope (Leica SP8X).

### Gene expression analysis

RNA was isolated using a RNeasy kit (Qiagen). Quantitative real-time polymerase chain reaction (qPCR) was performed on complementary DNA and quantified using the Pfaffl method.

### Traction Force Microscopy

Cells and fluorescent beads were imaged overnight. The reference image was obtained after removal of the cells. Timelapse images were aligned and cropped according to the reference image. Bead displacements were computed using Particle Image Velocimetry. Cell tractions were computed by Fourier transform-based traction microscopy. Traction magnitude was calculated as median value over space (the cell) followed by mean value over time.

### Z-Score analysis

The Z-score is a measure of a morpho-mechanical property *x* as a result of a specific environmental condition. The Z-score is defined as 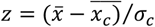 in which 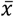 is the mean of morpho-mechanical property *x* within a specific culture condition, 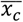 is the mean of physical property *x* of the control culture condition (E_P_-C_P_-WT) and *σ*_*c*_ is the standard deviation of physical property *x* of the control culture condition. Correlations between morpho-mechanical properties and culture conditions were calculated using cosine similarity analysis. An unsupervised clustering algorithm was used to detect clusters in the Z-score matrix.

### Statistics

Data is presented as mean ± standard error of the mean (SEM). Normality was tested using Shapiro Wilk test. Statistical tests used in this study were an unpaired student t-test (gene/protein expression analysis), a Mann-Whitney test and a Kruskall-Wallis test followed by Dunn’s multiple comparison test (quantification of morpho-mechanical state).

See additional details in SI Appendix, SI Materials and Methods

### Data and software availability

Software for the analysis of FAs (SFAlab) is publicly available through this [link](https://gitlab.tue.nl/stem/sfalab). Software for cosine similarity analysis and unsupervised clustering is publicly available through this [link](https://github.com/MinahilRaza/Bayesian-Inference-Criteria-based-Clustering).

## Acknowledgements

We thank John Eriksson (Åbo Akademi) for providing the VimWT and VimKO MEFs; Oscar Stassen and Rob Driessen (TU/e) for the help with the lentiviral system; Rob Hoeben and Martijn Rabelink (LUMC) for providing us with the vimentin shRNA and control plasmid; Emma Drabbe and Anouk van der Net (formerly of the TU/e) for the optimalisation of validation of the vimentin knockdown in ECs and VSMCs; Sylvia Dekker (TU/e) for the optimalisation of the western blot; Marta Sales-Pardo and Roger Guimerà (Universitat Rovira i Virgili) for the valuable discussion on the unsupervised clustering algorithm; Leon Hermans and Pim van den Bersselaar (TU/e, ICMS) for the fruitful discussions on experimental procedures and morpho-mechanical quantification. We gratefully acknowledge support by grants from the European Research Council (771168), the Netherlands Organization for Scientific Research (024.003.013), the Academy of Finland (307133, 316882, 330411 and 337531), the Åbo Akademi University Foundation’s Centers of Excellence in Cellular Mechanostasis (CellMech) and Bioelectronic Activation of Cell Functions (BACE) and EMJMD in Engineering of Data-Intensive Intelligent Software Systems (619819).

## Author Contributions

J.G., N.C.A.V.E., V.C., C.M.S. and C.V.C.B. designed the research. J.G. and N.C.A.V.E. performed experiments and analyzed the data. M.R. and S.A.R. contributed to the implementation of new analytic tools. V.C., C.M.S. and C.V.C.B. supervised the project. J.G., V.C., C.M.S. and C.V.C.B. wrote the paper. All authors reviewed the manuscript.

## Additional Information

**Supplementary Information** is available.

**Correspondence** and requests should be addressed to C.V., C.M.S. and/or C.V.C.B.

### Competing Interests

The authors declare no competing interests.

## Notes

### Competing Interest Statement

The authors have declared no competing interest.

## References

1. Vogel, V. & Sheetz, M. P. Mechanical forces matter in health and disease: from cancer to tissue engineering. In Nanotechnology (2010). doi:10.1002/9783527628155.nanotech057

2. Eichinger, J. F. et al.. Mechanical homeostasis in tissue equivalents: a review. Biomech. Model. Mechanobiol. 20, 833–850 (2021).

3. Cook, J. R. et al.. Abnormal muscle mechanosignaling triggers cardiomyopathy in mice with Marfan syndrome. J. Clin. Invest. 124, 1329–1339 (2014).

4. Cyron, C. J. & Humphrey, J. D. Vascular homeostasis and the concept of mechanobiological stability. Int. J. Eng. Sci. 85, 203–223 (2014).

5. Jaalouk, D. E. & Lammerding, J. Mechanotransduction gone awry. Nat. Rev. Mol. Cell Biol. 10, 63–73 (2009).

6. van Helvert, S., Storm, C. & Friedl, P. Mechanoreciprocity in cell migration. Nat. Cell Biol. 20, 8–20 (2018).

7. De Luca, M., Mandala, M. & Rose, G. Towards an understanding of the mechanoreciprocity process in adipocytes and its perturbation with aging. Mech. Ageing Dev. 197, 111522 (2021).

8. Humphrey, J. D. et al.. Mechanotransduction and extracellular matrix homeostasis. Nat. Rev. Mol. Cell Biol. 15, 802–812 (2014).

9. Miller, A. E., Hu, P. & Barker, T. H. Feeling Things Out: Bidirectional Signaling of the Cell–ECM Interface, Implications in the Mechanobiology of Cell Spreading, Migration, Proliferation, and Differentiation. Adv. Healthc. Mater. 9, 1–24 (2020).

10. Walker, M., Rizzuto, P., Godin, M. & Pelling, A. E. Structural and mechanical remodeling of the cytoskeleton maintains tensional homeostasis in 3D microtissues under acute dynamic stretch. Sci. Rep. 10, 1–16 (2020).

11. Ohashi, K., Fujiwara, S. & Mizuno, K. Roles of the cytoskeleton, cell adhesion and rho signalling in mechanosensing and mechanotransduction. J. Biochem. 161, 245–254 (2017).

12. Ingber, D. E. Tensegrity I. Cell structure and hierarchical systems biology. J. Cell Sci. 116, 1157–1173 (2003).

13. Ridge, K. M., Eriksson, J. E., Pekny, M. & Goldman, R. D. Roles of vimentin in health and disease. Genes Dev. 36, 391–407 (2022).

14. Steinert, P. M. & Roop, D. R. Molecular and cellular biology of intermediate filaments. Ann. Rev. Biochem. 57, 593–625 (1988).

15. Patteson, A. E. et al.. Vimentin protects cells against nuclear rupture and DNA damage during migration. J. Cell Biol. 218, 4079–4092 (2019).

16. Wu, H. et al.. Vimentin intermediate filaments and filamentous actin form unexpected interpenetrating networks that redefine the cell cortex. Proc. Natl. Acad. Sci. U. S. A. 119, 1–10 (2022).

17. Ivaska, J., Pallari, H. M., Nevo, J. & Eriksson, J. E. Novel functions of vimentin in cell adhesion, migration, and signaling. Exp. Cell Res. 313, 2050–2062 (2007).

18. Shemesh, T., Geiger, B., Bershadsky, A. D. & Kozlov, M. M. Focal adhesions as mechanosensors: A physical mechanism. Proc. Natl. Acad. Sci. U. S. A. 102, 12383–12388 (2005).

19. Mendez, M. G., Kojima, S. & Goldman, R. D. Vimentin induces changes in cell shape, motility, and adhesion during the epithelial to mesenchymal transition. FASEB J. 24, 1838–1851 (2010).

20. Gregor, M. et al.. Mechanosensing through focal adhesion-anchored intermediate filaments. FASEB J. 28, 715–729 (2014).

21. De Pascalis, C. et al.. Intermediate filaments control collective migration by restricting traction forces and sustaining cell-cell contacts. J. Cell Biol. 217, 3031–3044 (2018).

22. Choquet, D., Felsenfeld, D. P. & Sheetz, M. P. Extracellular matrix rigidity causes strengthening of integrin-cytoskeleton linkages. Cell 88, 39–48 (1997).

23. Langlois, B. et al.. Vimentin knockout results in increased expression of sub-endothelial basement membrane components and carotid stiffness in mice. Sci. Rep. 7, 1–15 (2017).

24. van Engeland, N. C. A. et al.. Vimentin regulates Notch signaling strength and arterial remodeling in response to hemodynamic stress. Sci. Rep. 9, 1–14 (2019).

25. Cox, T. R. & Erler, J. T. Remodeling and homeostasis of the extracellular matrix: Implications for fibrotic diseases and cancer. DMM Dis. Model. Mech. 4, 165–178 (2011).

26. Handorf, A. M., Zhou, Y., Halanski, M. A. & Li, W. J. Tissue stiffness dictates development, homeostasis, and disease progression. Organogenesis 11, 1–15 (2015).

27. Janmey, P. A., Hinz, B. & McCulloch, C. A. Physics and physiology of cell spreading in two and three dimensions. Am. Physiol. Soc. J. 36, 382–391 (2021).

28. Engler, A. et al.. Substrate Compliance Vs Ligand Density. Biophys. J. 86, 1–12 (2004).

29. Engler, A. J., Richert, L., Wong, J. Y., Picart, C. & Discher, D. E. Surface probe measurements of the elasticity of sectioned tissue, thin gels and polyelectrolyte multilayer films: Correlations between substrate stiffness and cell adhesion. Surf. Sci. 570, 142–154 (2004).

30. Zhang, Q., Yu, Y. & Zhao, H. The effect of matrix stiffness on biomechanical properties of chondrocytes. Acta Biochim. Biophys. Sin. (Shanghai). 48, 958–965 (2016).

31. Li, Y. et al.. Moving cell boundaries drive nuclear shaping during cell spreading. Biophys. J. 109, 670–686 (2015).

32. Wu, H. et al.. Vimentin intermediate filaments and filamentous actin form unexpected interpenetrating networks that redefine the cell cortex. Proc. Natl. Acad. Sci. U. S. A. 119, (2022).

33. Serra-Picamal, X., Conte, V., Sunyer, R., Muñoz, J. J. & Trepat, X. Mapping forces and kinematics during collective cell migration. Methods Cell Biol. 125, 309–330 (2015).

34. Wang, J. H.-C. Cell traction forces (CTFs) and CTF microscopy applications in musculoskeletal research. Oper Tech Orthop. 20, 106–109 (2010).

35. Jiu, Y. et al.. Vimentin intermediate filaments control actin stress fiber assembly through GEF-H1 and RhoA. J. Cell Sci. 130, 892–902 (2017).

36. Bazellières, E. et al.. Control of cell-cell forces and collective cell dynamics by the intercellular adhesome. Nat. Cell Biol. 17, 409–420 (2015).

37. Sales-Pardo, M., Guimerà, R., Moreira, A. A. & Nunes Amaral, L. A. Extracting the hierarchical organization of complex systems. PNAS 104, 15224–15229 (2007).

38. Calero-Cuenca, F. J., Janota, C. S. & Gomes, E. R. Dealing with the nucleus during cell migration. Curr. Opin. Cell Biol. 50, 35–41 (2018).

39. Peyton, S. R. & Putnam, A. J. Extracellular matrix rigidity governs smooth muscle cell motility in a biphasic fashion. J. Cell. Physiol. 204, 198–209 (2005).

40. Gaudet, C. et al.. Influence of Type I Collagen Surface Density on Fibroblast Spreading, Motility, and Contractility. Biophys. J. 85, 3329–3335 (2003).

41. Isomursu, A. et al.. Directed cell migration towards softer environments. Nat. Mater. 21, (2022).

42. Stroka, K. M. & Aranda-Espinoza, H. Neutrophils display biphasic relationship between migration and substrate stiffness. Cell Motil. Cytoskeleton 66, 328–341 (2009).

43. Dokukina, I. V. & Gracheva, M. E. A model of fibroblast motility on substrates with different rigidities. Biophys. J. 98, 2794–2803 (2010).

44. Harley, B. A. C. et al.. Microarchitecture of three-dimensional scaffolds influences cell migration behavior via junction interactions. Biophys. J. 95, 4013–4024 (2008).

45. Stanton, A. E., Tong, X., Lee, S. & Yang, F. Biochemical ligand density regulates Yes-associated protein translocation in stem cells through cytoskeletal tension and integrins.pd. ACS Appl. Mater. Interfaces 11, 8849–8857 (2019).

46. Vicente-Manzanares, M., Ma, X., Adelstein, R. S. & Horwitz, A. R. Non-muscle myosin II takes centre stage in cell adhesion and migration. Nat. Rev. Mol. Cell Biol. 10, 778–790 (2009).

47. Pelham, R. J. & Wang, Y. L. Cell locomotion and focal adhesions are regulated by substrate flexibility. Proc. Natl. Acad. Sci. U. S. A. 94, 13661–13665 (1997).

48. Rens, E. G. & Merks, R. M. H. Cell Shape and Durotaxis Explained from Cell-Extracellular Matrix Forces and Focal Adhesion Dynamics. iScience 23, 101488 (2020).

49. Ostrowska-Podhorodecka, Z. et al.. Vimentin tunes cell migration on collagen by controlling β1 integrin activation and clustering. J. Cell Sci. 134, 1–16 (2021).

50. Liu, C. Y., Lin, H. H., Tang, M. J. & Wang, Y. K. Vimentin contributes to epithelial-mesenchymal transition ancer cell mechanics by mediating cytoskeletal organization and focal adhesion maturation. Oncotarget 6, 15966–15983 (2015).

51. Swoger, M. et al.. Vimentin Intermediate Filaments Mediate Cell Morphology on Viscoelastic Substrates. ACS Appl. Bio Mater. 5, 552–561 (2022).

52. Colburn, Z. T. & Jones, J. C. R. Complexes of α6β4 integrin and vimentin act as signaling hubs to regulate epithelial cell migration. J. Cell Sci. 131, (2018).

53. Terriac, E. et al.. Vimentin Levels and Serine 71 Phosphorylation in the Control of Cell-Matrix Adhesions, Migration Speed, and Shape of Transformed Human Fibroblasts. Cells 6, (2017).

54. Venu, A. P. et al.. Vimentin supports directional cell migration by controlling focal adhesions. bioRxiv (2022).

55. Challa, A. A., Vukmirovic, M., Blackmon, J. & Stefanovic, B. Withaferin-A Reduces Type I Collagen Expression In Vitro and Inhibits Development of Myocardial Fibrosis In Vivo. PLoS One 7, e42989 (2012).

56. Challa, A. A. & Stefanovic, B. A Novel Role of Vimentin Filaments: Binding and Stabilization of Collagen mRNAs. Mol. Cell. Biol. 31, 3773–3789 (2011).

57. Ostrowska-podhorodecka, Z. et al.. Vimentin Regulates Collagen Remodeling Through Interaction with Myosin 10. Qeios [preprint] 1–28 (2022).

58. Levental, K. R. et al.. Matrix Crosslinking Forces Tumor Progression by Enhancing Integrin Signaling. Cell 139, 891–906 (2009).

59. PhotoCol User protocol. Advanced BioMatrix (2018). doi:10.1038/s41415-020-2365-1

