## Supplementary information for "Environmental stiffness restores mechanical homeostasis in vimentin-depleted cells"

[1]Department of Biomedical Engineering, Soft Tissue Engineering and Mechanobiology, Eindhoven University of Technology, Eindhoven, 5612AE, the Netherlands. [2]Institute for Complex Molecular Systems, Eindhoven University of Technology, Eindhoven, 5600MB, the Netherlands. [3]Faculty of Science and Engineering, Cell Biology, Åbo Akademi University, Turku, 20520, Finland. [4]Faculty of Science and Engineering, Information Technology, Åbo Akademi University, Turku, 20500, Finland. [5]Institute for Bioengineering of Catalonia, The Barcelona Institute of Science and Technology, Barcelona, 08036, Spain. Cecilia M. Sahlgren and Carlijn V.C. Bouten contributed equally to this work. Correspondence and requests for materials and codes should be addressed to V.C., C.M.S. and/or C.V.C.B. ( and/or).

**This PDF file includes:**

Supporting Materials and Methods
Supporting Figures S1 to S6
Supporting References

### Supporting Materials and Methods

**Cell culture.** *Mouse Embryonic Fibroblasts (MEFs)* - Vimentin wildtype (VimWT) and vimentin knockout (VimKO) MEFs (J. Eriksson, Åbo Akademi) were cultured in Advanced DMEM (Gibco) supplemented with 10% Fetal Bovine Serum (FBS, Greiner), 1% L-Glutamine (Lonza), and 1% penicillin/streptomycin (pen/strep, Lonza). *Human umbilical vein Endothelial Cells (ECs)* – ECs (Lonza) were cultured in Endothelial Basal Medium 2 (Promocell) supplemented 1% pen/strep (Lonza), 0.02 mg/ml fetal calf serum, 5 ng/ml epidermal growth factor, 10 ng/ml basic fibroblast growth factor, 20 ng/ml insulin-like growth factor, 0.5 ng/ml vascular endothelial growth factor 165, 1 µg/ml ascorbic acid, 22.5 µg/ml heparin and 0.2 µg/ml hydrocortisone (all supplements from Promocell). ECs were used up until passage number 5. During transduction, ECs were cultured in medium without pen/strep. *Human aortic Vascular Smooth Muscle Cells (VSMCs)* – VSMCs (Lonza) were cultured in growth medium consisting of medium 231 (Gibco) supplemented with 5% smooth muscle growth serum (Thermo Fisher) and 1% pen/strep. Cells cultured in growth medium are referred to as synthetic VSMCs. For differentiation towards contractile VSMCs, VSMCs were cultured in medium 231 supplemented with 1% smooth muscle differentiation serum (ThermoFisher) and 1% pen/strep for a minimum of 7 days. During transduction, VSMCs were cultured in medium without pen/strep. *Human Embryonal Kidney (HEK) 293T cells* - HEK 293T cells (Lonza) were cultured in advanced DMEM (Gibco) supplemented with heat-inactivated FBS and 1% pen/strep. For transfection, HEK 293T cells were cultured without pen/strep. All cells were cultured at 37°C and 5% $CO_2$, cells were passaged at confluency 80-90% and medium was replenished every 2-3 days.

***In vitro* vimentin knockdown of human ECs and VSMCs.** *(1) Lentiviral transfection* - One day prior to transfection, HEK 293T cells were seeded with 80% confluency. HEK 293T culture medium was refreshed with transfection medium several hours prior to transfection. HEK 293T cells were transfected with vimentin knockdown (VimKD) transfer plasmid pLKO.1-puro / VIM and non-targeting control (NTC) transfer plasmid pLKO.1-puro / SHC002. The transfection mix was prepared by mixing the transfer plasmid, envelope plasmid pMD2.G, packaging plasmid pCMVR8.74 mixed in polyethylenimine (PEI, Sigma Aldrich) in DMEM with a PEI/DNA ratio of 2:1. The transfection mix was dropwise added to the HEK 293T cells. After overnight incubation, medium was collected and refreshed thrice every 8 hours. The collected medium was centrifuged at 500g at 4°C for 5 minutes. The supernatant containing viral particles was collected, filtered with a 0.45µm filter unit and centrifuged at 50 000g at 4°C for 120 minutes. Supernatant was discarded and virus pellet was resuspended in sterile Phosphate Buffered Saline (PBS, Sigma), aliquoted and snapfreezed. *(2) Virus Titer Determination* - Virus titer determination was performed with HIV1 p24 SimpleStep ELISA kit (ab218268, Abcam). According to manufacturer's protocol, standard was prepared with p24 protein stock standard in sample diluent NS to make concentrations from 0 pg/mL - 300 pg/mL. Viral dilutions were made with NTC and VimKD aliquots in sample diluent NS. In short, the assay was performed by mixing standards and samples with detector and capture antibodies, incubated at room temperature (RT) for 1 hour on a plate shaker at 150 rpm. Mix was discarded and wells were washed three times with Wash Buffer PT. TMB substrate was incubated for 10 minutes on plate shaker at 300 rpm and the reaction was ended with Stop Solution. Absorbances were read at 450 nm in a plate reader. Results were used to interpolate sample data into standard curve to calculate virus concentration in pg/mL. To calculate TU/mL, the following equation was used: $Titer = (OD - b - blank)/m * 100 * dilution\ factor$, with absorbance (OD), intercept (b) and slope (m) of the standard curve. *(3) Lentiviral transduction* – ECs, synthetic VSMCs and contractile VSMCs were transduced with either VimKD virus or NTC virus as control. Transduction mix was made from transduction medium, supplemented with polybrene (14µL/mL, Sigma Aldrich) and virus (MOI = 0.25). ECs and VSMCs were incubated with the transduction mix at 37 °C and 5% CO2. Transduced ECs and VSMCs were selected by culturing the cells in growth medium supplemented with puromycin (1 µL/ml, Sigma Aldrich) for at least one week.

**Polyacrylamide (PAA) substrate preparation.** To facilitate binding of PAA gel to glass, glass bottom well plates (CellVis and MatTek) and microscope glass slides (ThermoFisher) were treated with a 14:1:1 solution of absolute ethanol (VWR), acetic acid (Merck) and Bind-Silane (Merck) respectively for 1h. Plates and slides were washed three times with absolute ethanol and dried using nitrogen. PAA gels with a Young's modulus of 12 kPa (physiological stiffness, $E_P$) and 27 kPa (high stiffness, $E_H$) were prepared in PBS according to the component mixtures indicated in Table 1. FluoSpheres were added to gels purposed for Traction Force Microscopy, while for all other gels latex beads were added. For 18mm gels (qPCR), 22µl PAA solution was pipetted on the treated glass. For 12mm gels (TFM & IF), 10µl PAA solution was pipetted on the treated glass. PAA droplet was covered with an 18 or 12 mm coverslip (VWR) and PAA gels were left to polymerize for 1h. After polymerization, PBS was added to the wells and coverslips were carefully removed. Gel stiffness was verified for all experiments with nano-indentation (Optics11 Piuma Nano-Indenter) using a probe with stiffness ~0.25N/m, tip radius ~25 µm, and indentation depth of 7 µm. The stiffness of each gel batch was determined from the average of 5 randomly independent indentations on the gel. Average of all measurements for each stiffness is displayed in Table S1.

| | **12 kPa ($E_P$)** | **27 kPa ($E_H$)** |
|---|---|---|
| *Measured stiffness ± SEM (in kPa)* | *13.5 ± 0.1* | *28.5 ± 0.3* |
| 40% acrylamide (Bio-Rad) | 18.8% | 26% |
| 2% bis-acrylamide (Bio-Rad) | 5% | 8% |
| FluoSpheres carboxylate modified microspheres (Invitrogen) or Carboxyl latex beads (Invitrogen) | 1% | 1% |
| 10% ammonium persulfate (Bio-Rad) | 0.5% | 0.5% |
| TEMED (Sigma-Aldrich) | 0.05% | 0.05% |

**Table S1.** PAA mixture concentrations.

**PAA gel culture***.* Before coating PAA gels, gels were functionalized using 1 mg/ml Sulfo-SANPAH (Pierce) for 5 minutes using 365nm wavelength ultraviolet light. Gels were washed three times 10 minutes in sterile PBS in a cell culture hood to remove the excess Sulfo-SANPAH. Gels were incubated overnight at 4°C with either 25µg/ml (physiological concentration, $C_P$) or 100µg/ml (high concentration, $C_H$) rat tail collagen type I (Corning) diluted in PBS. After overnight incubation, gels were washed twice with sterile PBS and incubated in medium for at least 20 minutes. Meanwhile, VimWT and VimKO cells were trypsinized using 0.25% trypsin-EDTA (Gibco) for approximately 6 minutes. For TFM and IF, 500 cells (in 50µl) were seeded per 12mm gel to have single cells. For qPCR, 100.000 cells (in 100µl) were seeded per 18mm gels to create a monolayer. Cells were left to attach for at least 4 hours before medium was added to the well. Cells were cultured for either 24h, 48h or 72h (specified in each figure).

**Protein expression analysis.** Protein expression levels were measured using western blotting. Samples were collected in Laemmlli buffer. After protein separation by SDS-PAGE in a 4-15% gradient gel (Bio-Rad), proteins were transferred to a nitrocellulose membrane during western blotting. Prior to incubation of primary antibodies, membranes were blocked with 5% bovine serum albumin (BSA, Roche) or non-fat dry milk in PBST for 30 minutes. Primary antibodies used in this study: ß-actin (4967L, Cell Signaling), Fibronectin (ab23751, abcam), HSC70 (ENZO life, 815-D) and Vimentin (ab20346, abcam). After incubation overnight at 4°C, membranes were washed three times with PBST and incubated with the secondary antibodies for 1 hour at room temperature. Samples were washed with PBST, incubated with the ECL western blotting substrate and visualized.

**Immunofluorescence staining.** Cells designated for IF were fixed using 4% paraformaldehyde for 15 minutes. Prior to and after fixation, cells were washed three times in PBS. Cells were permeabilized using 0.5% Triton-X-100 (Sigma) for 3 minutes and washed thrice with PBS-0.1% Tween20 (PBST, Sigma). Following permeabilization, samples were blocked with 3% BSA. Samples were incubated with primary antibody for 90 minutes at room temperature. Primary antibodies used in this study: Collagen type III (ab7778, abcam), Fibronectin (ab23751, abcam), Paxillin (ab32084, abcam), phosphorylated myosin light chain (pMLC; 3675S, Cell Signaling) and Vimentin (ab20346, abcam). After incubation with the primary antibody, samples were washed three times with PBST and incubated for 90 minutes with the secondary antibodies in PBST at room temperature. Some samples were also incubated with Phalloidin Atto 647 (Sigma) to stain F-actin and CNA probe (CNA35-OG488) to stain collagen. Then, samples were washed with PBST three times and incubated with 4'-6-diamidine-2'- phenylindole dihydrochloride (DAPI) for 15 minutes at RT, washed thrice and mounted with Mowiol. Stained samples were stored at 4°C. Images of ECM proteins and vimentin knockdown verification were obtained with a widefield epi-fluorescence microscope (Leica DMi8) using a 10x dry objective (0.32NA). Images of single cells stained for F-actin, pMLC and paxillin were obtained with a white light laser confocal microscope (Leica SP8X) using an oil-immersed 63x (1.4NA) objective. To quantify cell/nucleus height, the staining for actin/nucleus was also images in the X-Z and Y-Z plain through the center of the cell/nucleus.

**Gene expression analysis.** To analyze gene expression levels, quantitative real-time polymerase chain reaction (qPCR) was performed. Before collecting, cells were washed with PBST. RNA was isolated using a RNeasy kit (Qiagen) according to the manufacturer's protocol. Complementary DNA was made with dNTPs (Invitrogen), random hexadeoxynucleotides (Promega) and Moloney Murine Leukemia Virus Reverse Transcriptase (M-MLV; Invitrogen). qPCR was performed using SYBR Green (Biorad) and quantified by the Pfaffl method. Primers used are listed in Table S2.

| | Gene | Forward primer | Reverse primer |
|---|---|---|---|
| *MEF* | *Col3a1* | CACGTAAGCACTGGTGGACA | AGAAGTCTGAGGAATGCCAGC |
| | *Gapdh* | CAATGTGTCCGTCGTGGATCTG | TGTAGCCCAAGATGCCCTTCAG |
| | *Lox* | CGCAAAGAGTGAAGAACCA | GTGTCCTCCAGACAGAAGC |
| | *Mmp2* | AACTACGATGATGACCGGAAGTG | GGTCCTGAGAGTGTTCCAGCC |
| | *Timp2* | CCTGGGACACGCTTAGCATC | CATCCAGAGGCACTCATCCG |
| | *Vimentin* | GCTCCTGGATCTCTTCATCG | AAAGCACCCTGCAGTCATTC |
| *EC/VSMC* | *COL1A1* | AATCACCTGCGTACAGAACGG | TCGTCACAGATCACGTCATCG |
| | *COL3A1* | ATCTTGGTCAGTCCTATGC | TGGAATTTCTGGGTTGGG |
| | *FN1* | AAGACCAGCAGAGGCATAAGG | CACTCATCTCCAACGGCATAATG |
| | *GAPDH* | Primer Design | |
| | *VIMENTIN* | AAGACCTGCTCAATGTTAAGATC | CTGCTCTCCTCGCCTTCC |

**Table S2.** Genes used for qPCR with their forward and reverse primer pair.

**Traction Force Microscopy.** (1) *Time-lapse imaging* - Cells and beads were live imaged using a Leica DMi8 microscope using a 20x dry objective (0.55NA) and a 16-bit Hamamatsu Orca Flash sCMOS-camera equipped with temperature, CO2 and humidity control. Time-lapse imaging was performed for approximately 15 hours (overnight) with a time interval of 15 minutes and consisted of phase contrast imaging for visualization of the single cells and fluorescent imaging for visualization of the beads. After time-lapse imaging, cells were removed using several droplets of 5% sodium dodecyl sulfate (Sigma) diluted in water to acquire the reference image. *(2) Traction force microscopy analysis* - Before TFM analysis, images were aligned and cropped relative to the reference image. Bead displacements between any experimental time point and the reference image were computed using Particle Image Velocimetry by dividing the region of interest into smaller interrogation windows of 32 by 32 pixels with 0.5 overlap. The tractions were computed from the displacements by Fourier transform-based traction microscopy of an infinite gel with a finite gel thickness using the Boussinesq equation (1). *(3) Monolayer Stress Microscopy (MSM)* - MSM was used to compute stresses in single cells (2). In short, the tensorial stress state within the cell was computed as a mechanical equilibrium to the forces exerted by the elastic hydrogel to the cell which arise from the traction force of the cell on the hydrogel. Computations were executed in the custom 2D finite element method platform EMBRYO, a platform that was developed in the laboratory of J.J. Muñoz.

**Morphological and mechanical quantification in MATLAB.** *(1) Cell and nucleus morphology* - Nucleus/cell area was quantified by manually masking the nucleus or cell on a X-Y image using command *roipoly*. Using command *regionprops*, the number of pixels within the mask as a measure for nucleus/cell area and the number of pixels surrounding the mask as a measure for nucleus/cell perimeter were retrieved. Besides area, *regionprops* also fits an ellipse on top of the mask and returns the length of the long and short axis of this ellipse. Nucleus/cell shape was defined as the ratio between the short and long axis, meaning that a nucleus/cell is more elongated when cell shape is closer to 0 while a nucleus/cell is rounder when cell shape is closer to 1. Nucleus/cell height was quantified by manually measuring the distance between the bottom and top of the nucleus or cell on X-Z and Y-Z images using command *imtool*. Height quantification was performed by two persons to avoid personal bias. *(2) Cell traction force* – Traction magnitude of a cell at each timepoint was calculated by taking the median of all traction magnitude values within the mask surrounding the cell. Traction magnitude of a cell over time was calculated by taking the mean value over time. Every cell includes a minimum of 5 timepoints with a time interval of 30 minutes. Similar analysis holds for the calculation of average normal stress (ANS) and maximum shear stress (MSS). The ANS is defined as $\sigma_n = (\sigma_{11} + \sigma_{22})/2$ while the MSS is defined as $\sigma_s = (\sigma_{11} - \sigma_{22})/2$. *(3) Cell motility* – Migration pace and migration persistence were determined based on the centroid of the manual mask of the timelapse imaging. Migration pace is defined as the inverse of the length of the total path divided by the total time. Migration persistence is defined as the distance between start and end divided by the length of the total path. Similar as TFM analysis, a minimum of 5 timepoints was used. *(4) Focal adhesions (FAs)* - FAs were quantified using a customized MATLAB toolbox called SFA lab. This tool performs several pre-processing steps on a selected image followed by shape and orientation analysis on all identified objects within the pre-processed image. Pre-processing steps include top-hat filtering (*imtophat*), median filtering (*medilt2*), sharpen the greyscale (*imsharpen*), convert to binary image (*im2bw*) and select the size boundaries of the detected objects. The output of this tool per cell is a list of all FAs with information on the size and shape per FA. The number of FAs was determined by the length of the list. The FA size was determined by the mean of the size of all FAs of a cell. FA size fractioning was performed by setting fixed thresholds in size and count the number of FAs within these thresholds. The percentage was calculated by the number of FAs in between thresholds divides by the total number of FAs in a cell multiplied by 100%.

An average of these percentages over all cells were taken and plotted in a histogram. The total FA size was determined as the sum of the size of all FAs per cell. FA shape was quantified similarly as nucleus/cell morphology by dividing the short axis by the long axis of a fitted ellipse on top of a FA.

**Z-Score analysis.** *(1) Compute Z-score* - The Z-Score is a statistical tool to investigate the number of standard deviations the mean of a sample group is above or below the mean of the control group. This type of scoring is often used in high-throughput RNA interference screens to provide explicit information on the function of each siRNA relative to the rest (3). Within this study, the Z-Score is a measure of a morpho-mechanical property $x$ as a result of a specific environmental conditions. The Z-Score is defined as:

$$z = \frac{\bar{x} - \overline{x_c}}{\sigma_c}$$

in which $\bar{x}$ is the mean of morpho-mechanical property $x$ within a specific microenvironment, $\overline{x_c}$ is the mean of morpho-mechanical property $x$ of the control microenvironment ($E_P$-$C_P$-WT) and $\sigma_c$ is the standard deviation of morpho-mechanical property $x$ of the control microenvironment. *(2) Cosine similarity analysis* - In order to find correlations between morpho-mechanical properties and microenvironments, we calculated the cosine similarity between vectors within the Z-score matrix. For clarity, a Z-score matrix with $C$ rows (microenvironment condition) and $P$ columns (morpho-mechanical properties) would yield two cosine similarity matrices with dimensions $C$x$C$ and $P$x$P$ respectively. Each element of the similarity matrix was computed using the dot product of vectors. The normalized dot product between the $i^{\text{th}}$ and $j^{\text{th}}$ row $C$ would yield the element $A_{ij}$ and was calculated using the following equation:

$$A_{ij} = \frac{\vec{C_i} \cdot \vec{C_j}}{|\vec{C_i}||\vec{C_j}|}$$

This was repeated over all rows and columns creating one similarity matrix of environmental conditions and one similarity matrix of morpho-mechanical properties. If $A_{ij}$ is close to 1, this means that the two environmental conditions or morpho-mechanical properties have a high correlation. If $A_{ij}$ is close to -1, this means that the two culture conditions or physical properties behave very different from each other. *(3) Unsupervised clustering* – Next, we ordered the two similarity matrices such that the variables with higher correlation were placed closer to the diagonal and the ones with lower correlation were close to the matrix edges. We used simulated annealing to order a similarity matrix of size $N$x$N$ using the cost function $C$ described as:

$$C = {}^{1}/_{N} \sum_{i,j=1}^{N} A_{ij}|i - j|$$

Each iteration consisted of 10 steps. For every step, we randomly picked a segment of $W$ contiguous rows (where $W$ is chosen from Gaussian distributions whose variance is a function of temperature $T$ and size $N$ of the matrix) and rearranged the first row based on a uniform random distribution. The remaining rows were arranged in a way that their relative distance to this row remains similar. We computed the cost function $C'$ for this order. If the cost decreased, we accepted the order. Even if the cost did not decrease, we accepted the order with a probability indicated as:

$$p = e^{\frac{C-C'}{T}}.$$

The temperature was decreased in every iteration by a factor of 0.95. Note that during the start of the annealing process, the probability of accepting an order which did not decrease the cost function is higher.

This is called the exploration phase. As the temperature cools down, this probability decreased. This phase is referred to as exploitation. Simulated annealing decreases the cost function by exploring the search space without getting stuck in local minima of the cost function *C*. The iterations stop when *C* has not changed for 20 iterations. Once the matrix has been ordered, we used a greedy approach to find the most optimal partition based on the Bayesian Inference Criterion (BIC). The best partition is the one which minimizes the BIC score:

$$BIC = m \cdot \ln\left(\frac{\mathcal{L}}{m}\right) + k \ln(m)$$

where $k$ is the number of parameters in the model, $\mathcal{L}$ is the sum of least squares (residuals) for the partition and $m$ is the number of datapoints.

**Statistics.** Data is presented as mean ± standard error of the mean (SEM). Statistical analyses were performed using Graphpad Prism. Normality was tested using the Shapiro-Wilk test. For gene and protein expression analysis, data passed the normally test and significant differences between two groups were tested using an unpaired student t-test. Data on the morphological and mechanical state did not pass the normality test and was analyzed using non-parametric tests. Significance between different substrate conditions within each cell type was tested using the Kruskall-Wallis test followed by Dunn's multiple comparison test. Significance between cell types on a specific substrate was tested using the Mann-Whitney test. Significant differences between different substrate conditions of VimWT cells are displayed in green while significant differences between different substrate conditions of VimKO cells are displayed in red. Significant differences between VimWT cells and VimKO cells are displayed in black. Denotations: **** $p<0.0001$; *** $p<0.001$; ** $p<0.01$; * $p<0.05$; · $p<0.1$.

### Supporting Figures

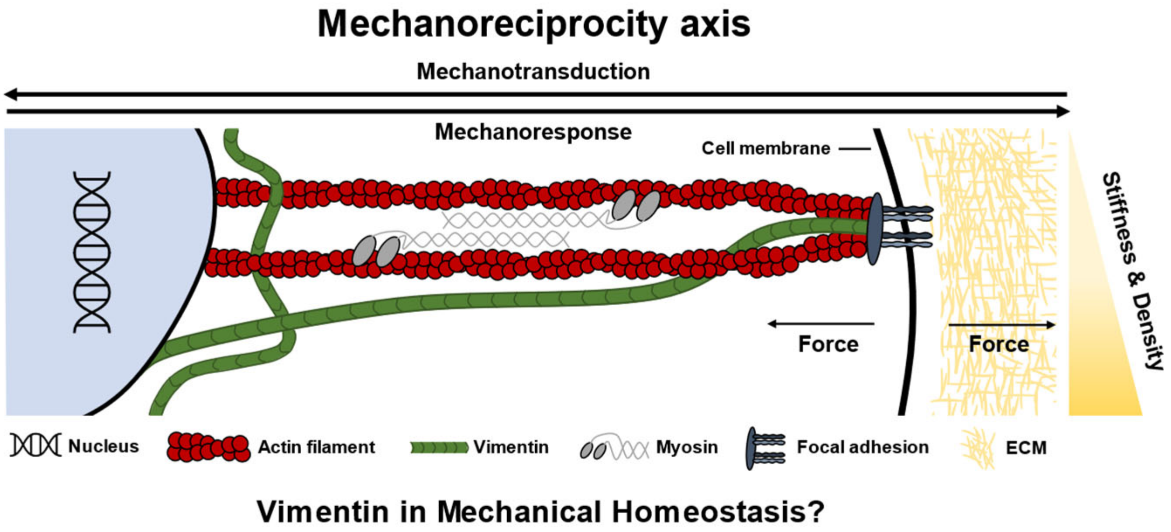


**Figure S1. Schematic representation of mechanoreciprocity in mechanical homeostasis.** Cells can sense mechanical signals (e.g. matrix stiffness and density) from its microenvironment and covert these into biomechanical and biochemical signals in the nucleus. These signals trigger a response in which the cell adapt is morphological and mechanical (morpho-mechanical) state of the cell as well as synthesis and remodelling of the ECM. This dynamic bi-directional mechanical interplay between the cell and its microenvironment is defined as mechanoreciprocity. This process is crucial for cells and tissues to establish, maintain and restore a preferred morpho-mechanical state and therefore mechanical homeostasis. Intermediate filament vimentin has been shown to associate with all single components along the mechanoreciprocity axis, however, how vimentin contributes to mechanical homeostasis is yet to be discovered.

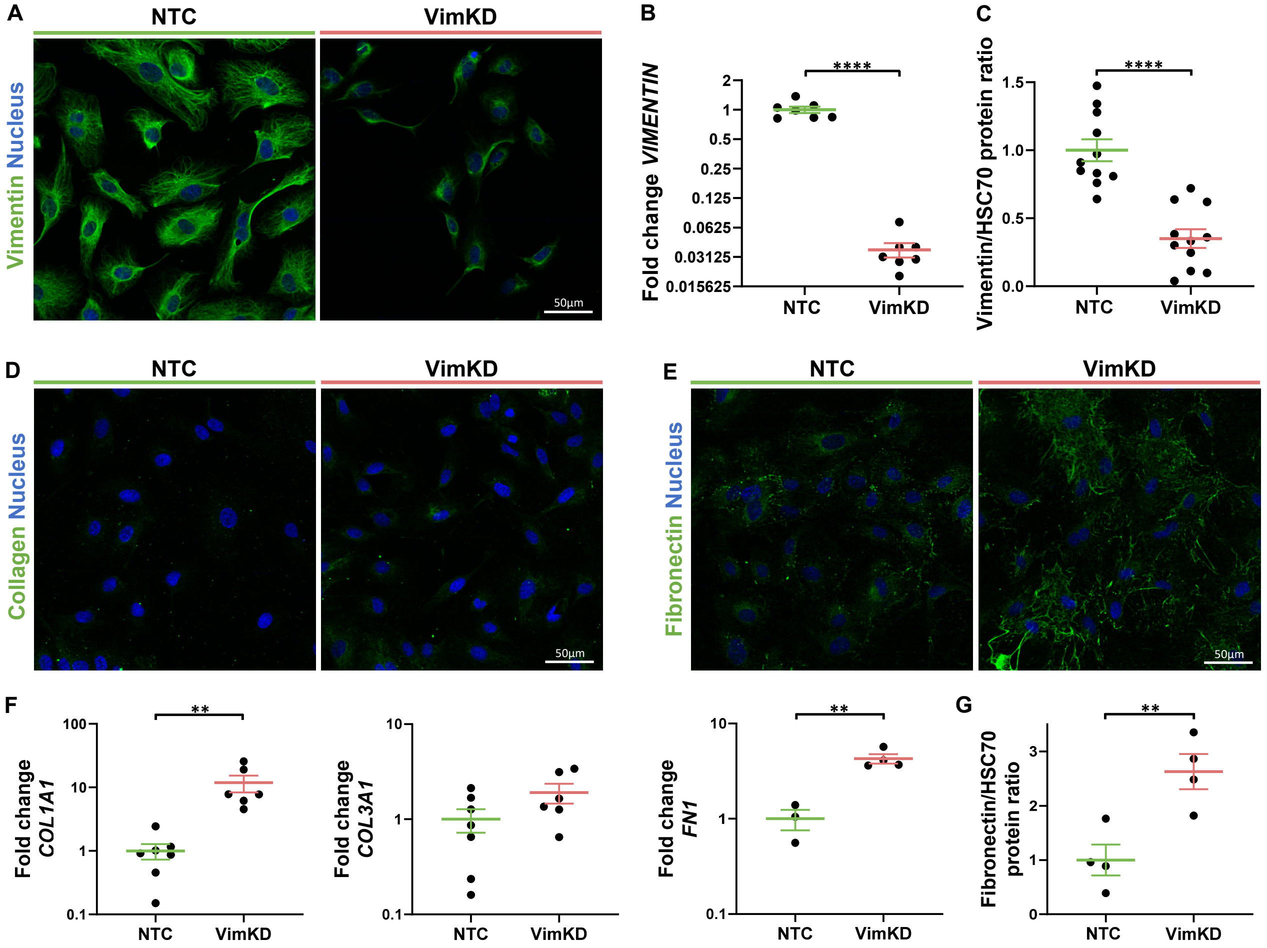


**Figure S2. Vimentin depletion increases ECM production *in vitro* in endothelial cells.** (A-C) Vimentin knockdown verification in endothelial cells on vimentin presence by immunofluorescence staining for vimentin (A), vimentin gene expression by q-PCR, with respect to reference gene *GAPDH*, normalized to NTC (B) and vimentin protein quantification by western blotting, normalized to NTC (C). (D-G) ECM expression in VimWT and VimKD endothelial cells on protein level by immunofluorescence staining for collagen (D) and fibronectin (E), on gene level by q-PCR for ECM genes *COL1A1*, *COL3A1* and *FN1,* with respect to reference gene *GAPDH*, normalized to NTC (F) and fibronectin protein quantification by western blotting, normalized to NTC (G). Data is represented as mean±SEM. An unpaired t-test was used for statistical analysis. Denotations: **** p<0.0001; *** p<0.001; ** p<0.01; * p<0.05; · p<0.1.

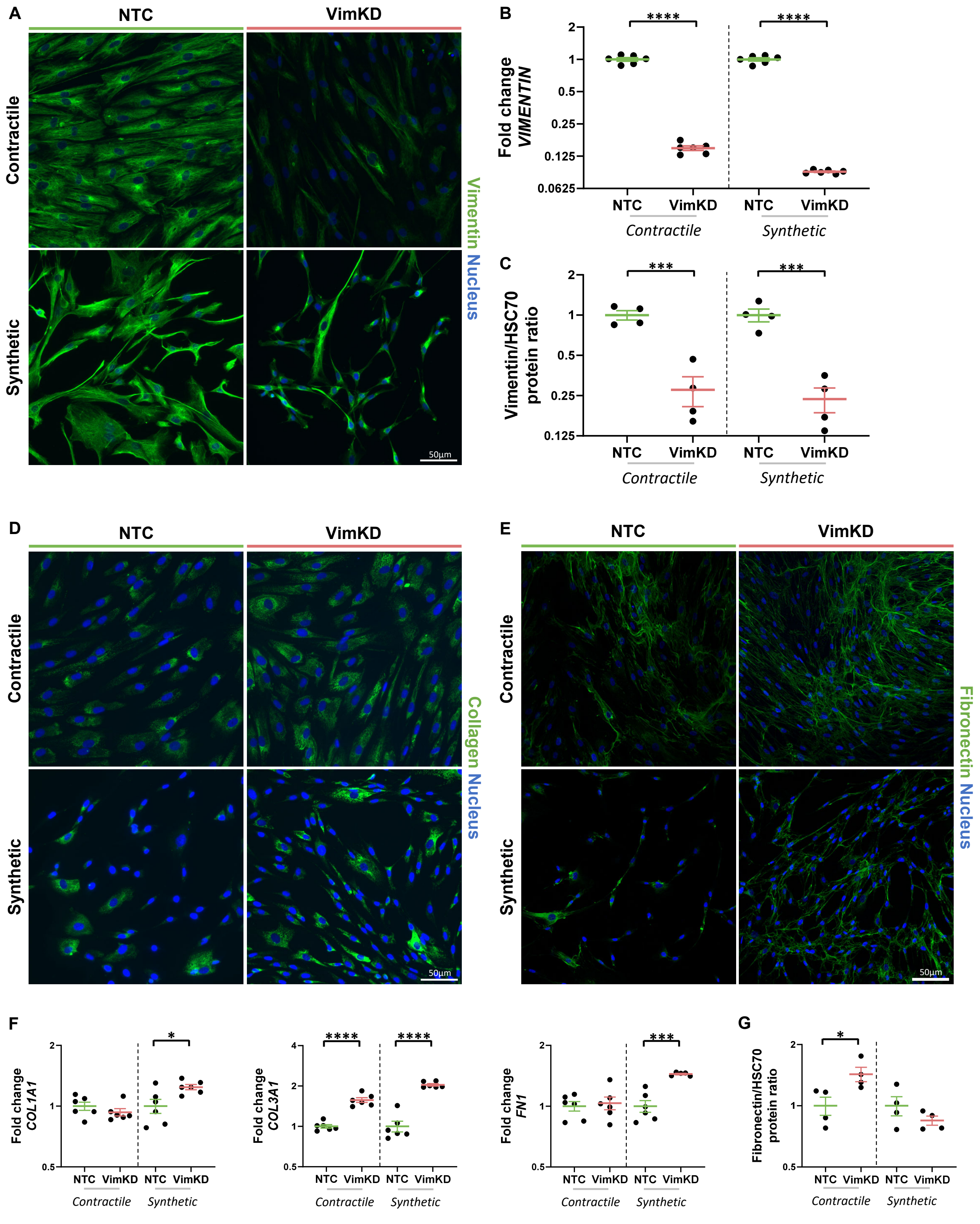


**Figure S3. Vimentin depletion increases ECM production *in vitro* in vascular smooth muscle cells.** (A-C) Vimentin knockdown verification in vascular smooth muscle cells on vimentin presence by immunofluorescence staining for vimentin (A), vimentin gene expression by q-PCR, with respect to reference gene *GAPDH*, normalized to NTC per phenotype (B) and vimentin protein quantification by western blotting, normalized to NTC per phenotype (C). (D-G) ECM expression in VimWT and VimKD vascular smooth muscle cells on protein level by immunofluorescence staining for collagen (D) and fibronectin (E), on gene level by q-PCR for ECM genes *COL1A1*, *COL3A1* and *FN1*, with respect to reference gene *GAPDH*, normalized to NTC per phenotype (F) and fibronectin protein quantification by western blotting, normalized to NTC per phenotype (G). Data is represented as mean±SEM. An unpaired t-test was used for statistical analysis. Denotations: **** $p<0.0001$; *** $p<0.001$; ** $p<0.01$; * $p<0.05$; · $p<0.1$.

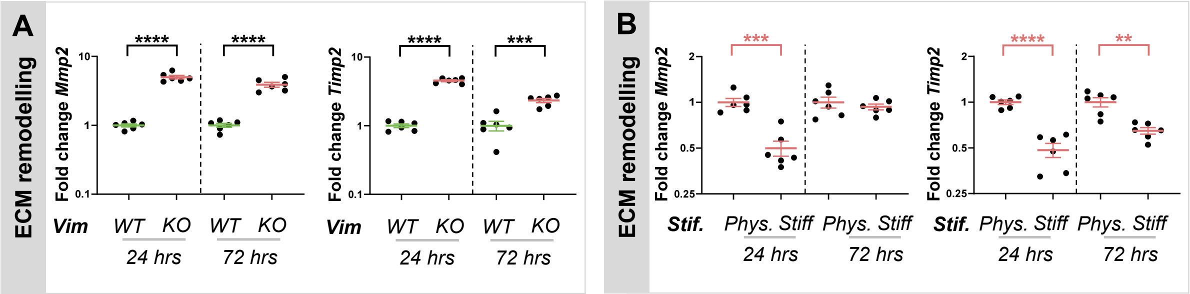


**Figure S4. Increased substrate stiffness compensates for vimentin depletion altered ECM remodelling.** Gene expression after 24 and 72 hours by q-PCR of ECM remodelling genes *Mmp2* and *Timp2* with respect to *Gapdh* expression. (A) VimWT and VimKO cells cultured on PAA gels with physiological substrate stiffness (12 kPa), VimKO expression levels normalized to VimWT expression levels. (B) VimKO cells cultured on PAA gels with physiological substrate stiffness (Phys.) and on a stiff substrate (glass), stiff substrate expression levels normalized to physiological expression levels. N=6. Data is represented as mean±SEM. An unpaired t-test was used for statistical analysis. Denotations: **** $p<0.0001$; *** $p<0.001$; ** $p<0.01$; * $p<0.05$; · $p<0.1$.

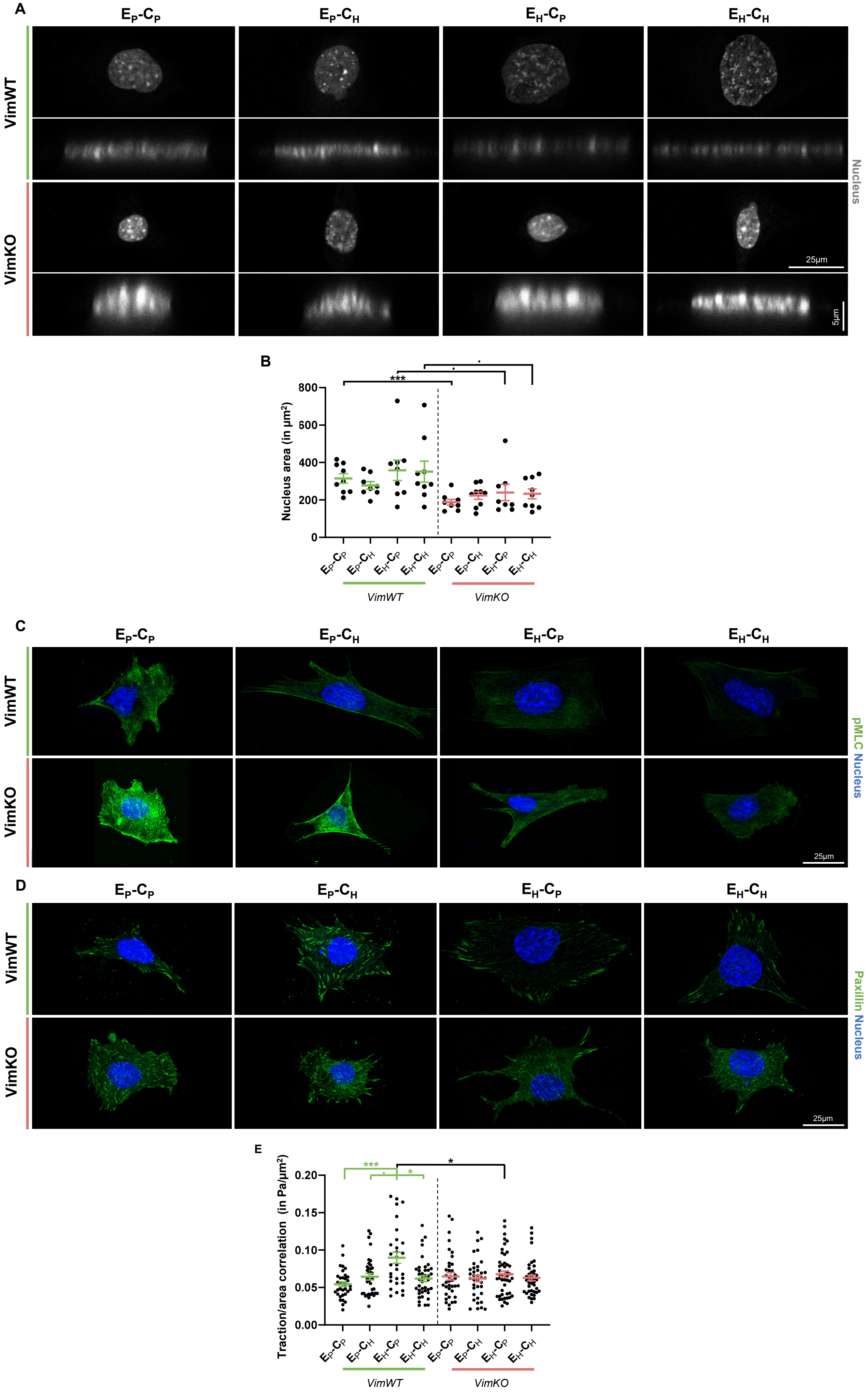

A
$E_P$-$C_P$
$E_P$-$C_H$
$E_H$-$C_P$
$E_H$-$C_H$
VimWT
VimKO
Nucleus
25µm
5µm
B
Nucleus area (in µm²)
VimWT
VimKO
C
pMLC Nucleus
D
Paxillin Nucleus
E
Traction/area correlation (in Pa/µm²)

**Figure S5. Vimentin-depleted cells have a differential morpho-mechanical phenotype of the cell.** (A) Top and side confocal images of VimWT and VimKO cells cultured on different substrate conditions for 24 hours stained with DAPI for nucleus visualization. (B) Nucleus area quantification was based on top images of nucleus based on immunostaining for DAPI (images displayed in panel A). Each dot represents one cell. (C) Traction area correlation was calculated by dividing traction magnitude by area of the cell. Each dot represents one cell over time (within timerange 12-24 hours). (D-E) Immunofluorescence staining of VimWT and VimKO cells on different substrates for phosphorylated myosin light chain (pMLC) as a measure for cell tension (D) and paxillin as a measure for focal adhesions (E). Data is represented as mean±SEM. Kruskall-Wallis followed by Dunn's multiple comparison was used for statistical analysis between substrate conditions within VimWT cells (displayed in green) and VimKO cells (displayed in red). An unpaired t-test was used for statistical analysis between VimWT and VimKO cells on different substrate conditions (displayed in black). Denotations: **** $p<0.0001$; *** $p<0.001$; ** $p<0.01$; * $p<0.05$; · $p<0.1$.

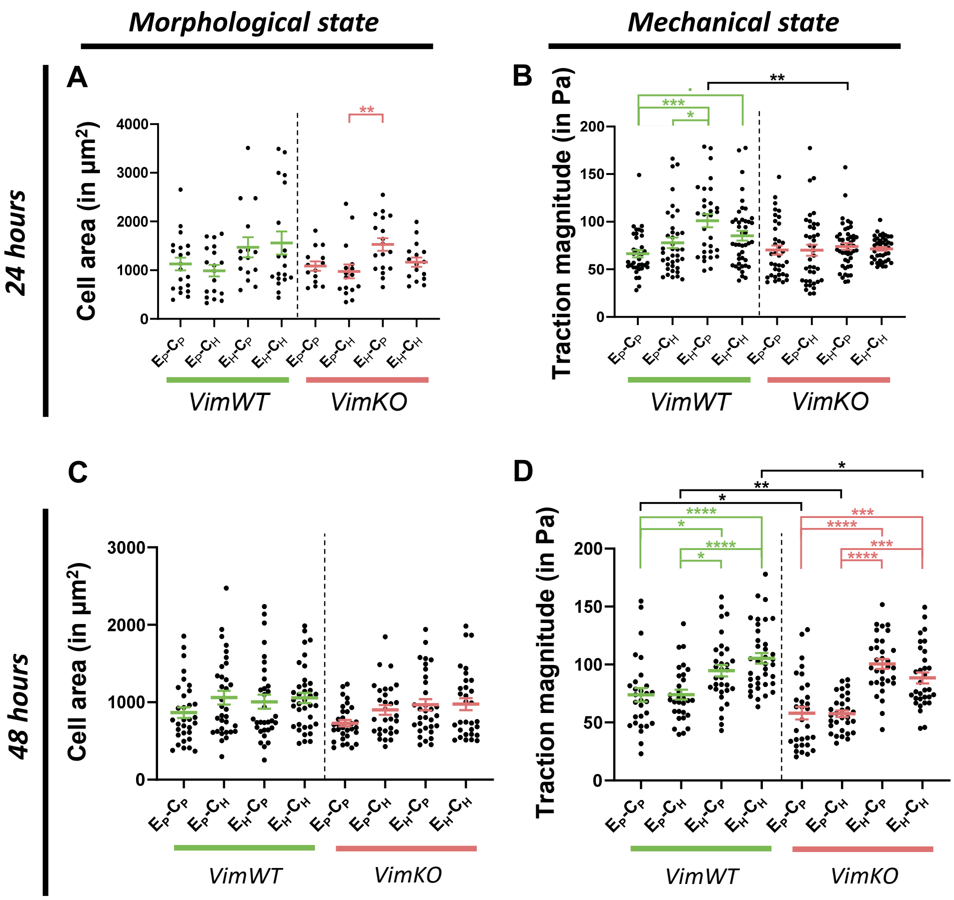


**Figure S6. Vimentin-depleted cells have the ability to respond in a similar manner as vimentin-presenting cells to changes in the microenvironment after 48 hours.** VimWT and VimKO cells were cultured on different environmental conditions for 24 hours (A-B) and results were compared to 48 hours (C-D). Morphological state was quantified by cell area (A&C) and mechanical state was quantified by traction force magnitude (B&D), (A&C) Cell area was quantified based on the mask of immunofluorescence (IF) staining for F-actin (A; n=15+) or the mask of phase contrast images (C; n=31+). Each dot presents one cell. (B&D) Traction magnitude was measured with Traction Force Microscopy. Each dot represents on cell over time within timerange 12-24 hours (B) or 36-48 hours (D) and is calculated as median over spatial map of the cell and mean of time (N=31+). Data is represented as mean±SEM. Kruskall-Wallis followed by Dunn's multiple comparison was used for statistical analysis between substrate conditions within VimWT cells (displayed in green) and VimKO cells (displayed in red). A Mann-Whitney test was used for statistical analysis between VimWT and VimKO cells on different substrate conditions (displayed in black). Denotations: **** $p<0.0001$; *** $p<0.001$; ** $p<0.01$; * $p<0.05$; · $p<0.1$.

### Supporting References


1. X. Trepat, *et al.*, Physical forces during collective cell migration - supplementary information. *Nat. Phys.* **5**, 426–430 (2009).
2. X. Serra-Picamal, V. Conte, R. Sunyer, J. J. Muñoz, X. Trepat, Mapping forces and kinematics during collective cell migration. *Methods Cell Biol.* **125**, 309–330 (2015).
3. A. Birmingham, *et al.*, Statistical methods for analysis of high-throughput RNA interference screens. *Nat. Methods* **6**, 569–575 (2009).
